# The LudP arrest peptide stalls translation by remodeling the ribosomal A-site

**DOI:** 10.64898/2026.10.26.761930

**Authors:** Felix Gersteuer, Yuki Harada, Keigo Fujiwara, Haaris A. Safdari, Alena Harreither, Lars V. Bock, Daniel N. Wilson, Shinobu Chiba

**Affiliations:** Institute for Biochemistry and Molecular Biology, Martin-Luther-King-Platz 6, University of Hamburg, 20146 Hamburg, Germany; Faculty of Life Sciences and Institute for Protein Dynamics, Kyoto Sangyo University, Kamigamo, Motoyama, Kita-ku, Kyoto 603-8555, Japan; Department of Gene Function and Phenomics, National Institute of Genetics, Mishima, Japan; Department of Theoretical and Computational Biophysics, Max Planck Institute for Multidisciplinary Sciences, Göttingen, Germany

## Abstract

Bacterial ribosome arrest peptides couple translation to secretion and membrane homeostasis, providing an intracellular regulatory feedback system. Although several Sec/YidC-associated arrest peptides have been characterized, how evolutionarily diverse peptides converge on equivalent physiological outcomes remains unclear. Here we identify LudP (**L**acrimispora **u**pstream of *sec**D**F* peptide), a previously unrecognized arrest peptide conserved in the genus *Lacrimispora* and encoded immediately upstream of *secDF*. In vitro and in vivo assays show that LudP induces robust Gram-positive– specific translation arrest that is released in a Sec signal–dependent manner. Using systematic mutagenesis and deep mutational scanning, we define a compact 15-residue C-terminal arrest element that governs arrest stability and A-site residue selectivity. Cryo-electron microscopy reveals that a LudP nascent-chain α-helix in the exit tunnel remodels the peptidyltransferase center, with Asp133-dependent conformational rearrangements of 23S rRNA, notably U2506, sterically and energetically destabilizing productive A-site aminoacyl–tRNA accommodation. These findings support functional convergent evolution of secretion-coupled arrest peptides across bacterial lineages.

## Introduction

Ribosome arrest peptides monitor the intracellular environment through translation arrest and thereby contribute to cellular homeostasis by feedback regulation of gene expression^1–4^. In bacteria, several arrest peptides have been identified that maintain the homeostasis of protein localization pathways, including SecM from *Escherichia coli*^5^, MifM from *Bacillus subtilis*^6^, VemP from *Vibrio alginolyticus*^7^, and CliM from members of the class *Clostridia*^8^. Among these, *E. coli* SecM regulates the intracellular level of SecA, the ATPase that drives the Sec protein secretion pathway^5^. Under normal conditions, expression of *secA* is repressed because its Shine–Dalgarno (SD) sequence is sequestered within an mRNA stem-loop structure. When translation arrest occurs near the 3′ end of *secM*, however, the stalled ribosome maintains the stem-loop in an unwound conformation, thereby continuously exposing the SD sequence and allowing sustained translation of *secA*. SecM is a 170-amino-acid secretory protein containing an N-terminal Sec signal peptide and a C-terminal arrest motif (FXXXXWIXXXXGIRAGP) that specifically interacts with the ribosome to induce translation arrest^9,10^. As SecM is translocated through the Sec translocon, the nascent chain is pulled from its N-terminus, disrupting the interaction between the arrest motif and the ribosome and thereby releasing the stalled complex. Therefore, SecM-mediated arrest persists when Sec translocation is impaired, enabling *E. coli* to induce SecA synthesis in response to reduced secretion capacity^5^. VemP employs a similar mechanism to regulate expression of the Sec component SecDF^7^, whereas MifM and CliM feedback-regulate expression of the membrane protein insertase YidC^6,8^. In all cases, these arrest peptides are encoded immediately upstream of the genes whose expression they regulate.

To comprehensively identify previously unrecognized arrest peptides involved in maintaining the homeostasis of the Sec and YidC pathways, we recently searched more than 30,000 bacterial genomes for genes sharing characteristic features of known arrest peptides^11,12^. Specifically, we focused on small secretory or membrane proteins encoded upstream of *sec* or *yidC* genes that contained highly conserved motifs near their C-termini, and examined their arrest activities using both *in vivo* and *in vitro* assays. This screen identified nearly twenty arrest peptides encoded upstream of *sec* or *yidC* genes^11,12^. Like SecM and MifM, these peptides are presumed to regulate expression of downstream *sec* or *yidC* genes through translation arrest; however, experimental evidence supporting this physiological role remains unavailable for most of these candidates.

In the present study, we re-examined the genomic dataset generated in our previous screen to identify additional factors that may have been overlooked despite sharing common features with known arrest peptides. This analysis identified a previously unrecognized arrest-peptide candidate in the genus *Lacrimispora* within the class *Clostridia*, which is encoded immediately upstream of *secDF*. We therefore investigated whether this protein, which we named LudP (for **L**acrimispora **u**pstream of sec**D**F **p**eptide), functions as a ribosome arrest peptide. We demonstrate that LudP induces translation arrest on *B. subtilis* ribosomes and that this arrest is released in a Sec signal-dependent manner, a hallmark shared by previously characterized arrest peptides that monitor the Sec pathway. To elucidate the molecular basis of LudP-mediated translation arrest, we combined comprehensive mutational analyses with cryo-electron microscopy. These analyses revealed that LudP induces ribosome stalling through interactions between a 14-residue C-terminal arrest motif and the ribosome. We further identified the residues essential for arrest and defined the molecular interactions responsible for this activity. Moreover, our cryo-EM structure, together with molecular dynamics (MD) simulations, show that LudP’s DGEK motif in cooperation with a N-terminal helix inhibits stable accommodation of the A-site aminoacyl-tRNA by displacing nucleotide U2506 of 23S rRNA toward the A-site. Ultimately, our findings define LudP as a distinct mechanistic archetype of secretion-monitoring peptides, providing concrete structural evidence of how diverse bacterial lineages have independently evolved unique chemical toolkits to co-opt the exact same vulnerable sites within the ribosomal peptidyltransferase center (PTC).

## Results

### LudP induces ribosome stalling on *B. subtilis* ribosomes *in vitro*

In our previous study, we identified nearly twenty arrest peptides by comprehensively screening upstream open reading frames (uORFs) associated with *sec* genes^12^. Upon further careful examination of the genomic dataset generated in that study, we identified several additional genes sharing characteristic features of known arrest peptides. One of them was LudP (**L**acrimispora **u**pstream of *sec**D**F* **p**eptide). LudP homologs were present in a subset of bacteria belonging to the class *Clostridia* (**Supplementary Fig. 1a**). In this study, we focused on the LudP homolog from *Lacrimispora xylanolytica*. *L. xylanolytica* LudP is a small 147-amino-acid protein encoded immediately upstream of *secDF* and contains a putative N-terminal signal peptide. A highly conserved sequence near its C terminus (residues 129–137; IHNSDGEKE) is well conserved among LudP homologs (consensus sequence: I-H-D/N-S-D-G-E/K-K-G/E) (**Supplementary Fig. 1b,c**).

To determine whether *L. xylanolytica* LudP induces ribosome stalling, we analyzed its translation using the PURE system, a reconstituted *in vitro* translation system consisting of purified *E. coli* translation components together with T7 RNA polymerase for transcription^13^. In addition to the original *E. coli*-based PURE system (hereafter Ec PURE), we employed a hybrid *B. subtilis* PURE system (hereafter Bs PURE), in which purified *B. subtilis* ribosomes replace *E. coli* ribosomes^14^. As an *in vitro* arrest reporter, we constructed a translational fusion in which the C-terminal region of LudP (residues 16–147) lacking the intact signal peptide was inserted between GFP and LacZα, encoding the N-terminal ∼60 amino acids of LacZ (**Fig. 1a**). The reporter was transcribed and translated in both Ec PURE and Bs PURE, and the translation products were analyzed by Western blotting. A D133A mutant, in which the highly conserved Asp133 was replaced with Ala, was analyzed in parallel.

**Fig. 1.**
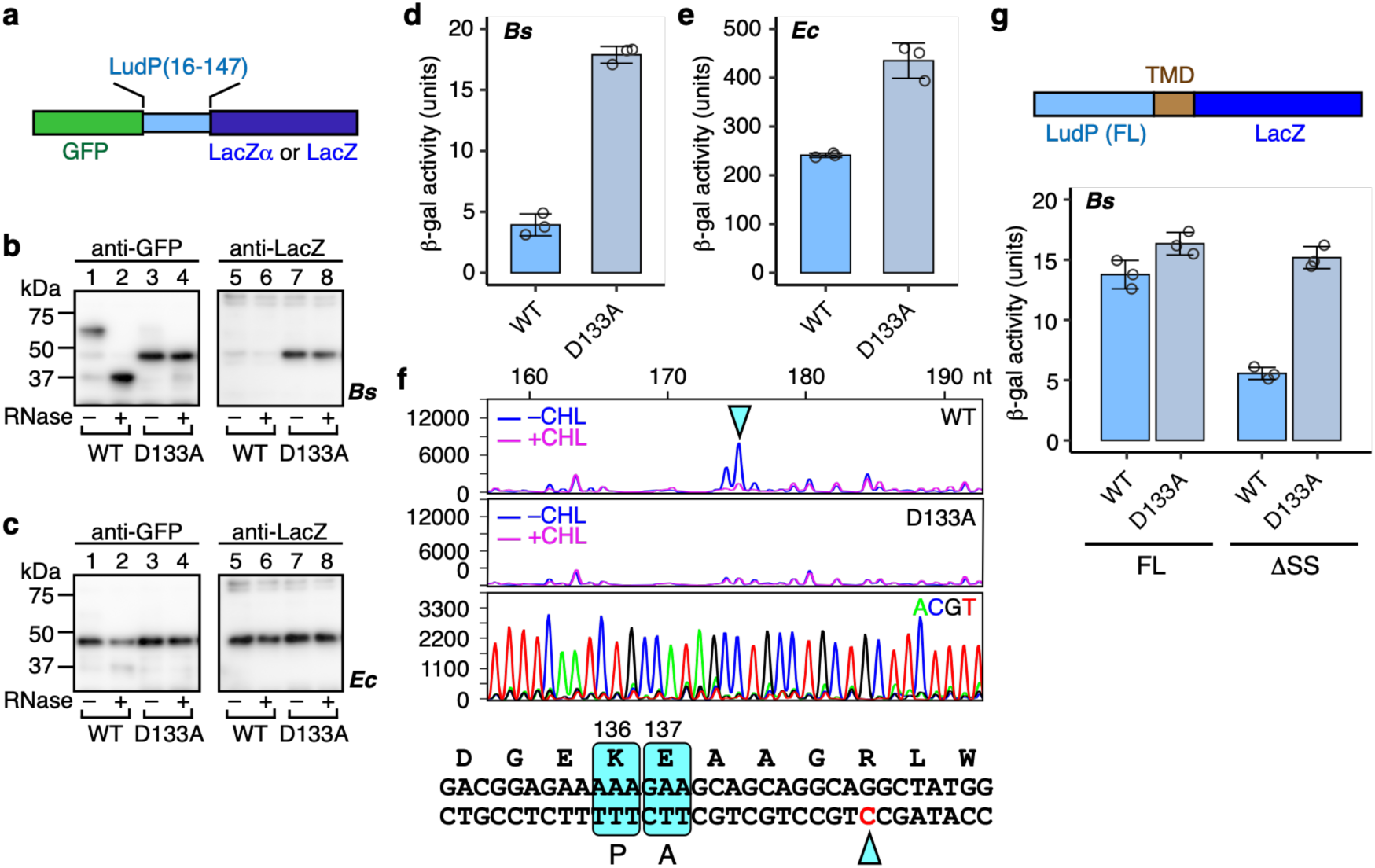
LudP induces translation arrest in *B. subtilis*. **a**. Schematic representation of the *gfp-ludP(16-147)-lacZα* and *gfp-ludP(16-147)-lacZ* reporters used for in vitro and in vivo experiments, respectively. **b, c.** *In vitro* translation assay. *gfp-ludP(16-147)-lacZα* was translated in Bs PURE (b) or Ec PURE (c) at 37°C for 20 min. Translation products were analyzed by immunoblotting with anti-GFP (left) and anti-LacZ (right) antibodies. A portion of each sample was treated with RNase A to digest the tRNA moiety of peptidyl-tRNA. **d, e**. β-galactosidase activity (mean ± s.d., n = 3, biologically independent cultures) of *B. subtilis* (d) or *E. coli* (e) cells harboring the wild-type (WT) or D133A derivative of the *gfp-ludP(16-147)-lacZ* reporter. **f.** Toeprinting analysis of WT (upper) and D133A (middle) LudP reporters translated in Bs PURE. Reverse transcription products were analyzed by a capillary electrophoresis. *In vitro* translation was performed in the presence or absence of chloramphenicol (CHL). Toeprint length (nt) was calibrated against dideoxy sequencing products (lower; ACGT). P- and A-site codons, inferred from the size of the toeprint product (cyan triangle), are indicated as P and A, respectively, together with their codon numbers (bottom). **g**. Schematic representation of the *ludP(FL)-TMD-lacZ* reporter and β-galactosidase activity (mean ± s.d., n = 3, biologically independent cultures) of *B. subtilis* cells harboring the wild-type (WT) or D133A derivative of the *ludP(FL)-TMD-lacZ* reporter. FL and ΔSS indicate the full-length and signal-sequence-deleted variants of *ludP*, respectively.

Translation of the wild-type reporter in Bs PURE produced a major band migrating between the 50-and 75-kDa molecular weight markers (**Fig. 1b**, lane 1). This product was detected only by the anti-GFP antibody, suggesting that it lacks the C-terminal LacZα moiety. RNase treatment induced a mobility shift of this product (**Fig. 1b**, lanes 1 and 2), indicating that it represents a stalled peptidyl-tRNA. In contrast, translation of the D133A mutant generated a major product migrating between the 37-and 50-kDa markers whose mobility was unaffected by RNase treatment and which was also detected with an anti-LacZ antibody, indicating that it corresponds to the full-length translation product. When the same constructs were translated in Ec PURE, both the wild-type and D133A reporters predominantly produced full-length products (**Fig. 1c**). These results demonstrate that *L. xylanolytica* LudP induces robust stalling on *B. subtilis* ribosomes, but not on *E. coli* ribosomes.

We next examined the LudP homolog from *Lacrimispora indolis*. Similar to *L. xylanolytica* LudP, the *L. indolis* homolog accumulated predominantly as a stalled product in Bs PURE and, unlike *L. xylanolytica* LudP, also accumulated predominantly as the stalled product in Ec PURE (**Supplementary Fig. 2**). Nevertheless, these results indicate that LudP homologs from both *L. xylanolytica* and *L. indolis* efficiently stall *B. subtilis* ribosomes, consistent with the limited phylogenetic distribution of LudP in Gram-positive bacteria (**Supplementary Fig. 1a**).

### LudP induces translation arrest *in vivo*

To determine whether LudP also induces translation arrest *in vivo*, we constructed a translational fusion in which the C-terminal region of *L. xylanolytica* LudP (residues 16– 147) was inserted between GFP and full-length LacZ (**Fig. 1a**). The reporter was expressed in *B. subtilis* and *E. coli*, and translation arrest was evaluated by measuring β-galactosidase activity. Efficient translation arrest within the LudP region prevents translation from reaching the downstream *lacZ* gene, resulting in low β-galactosidase activity.

In *B. subtilis*, the wild-type reporter exhibited 3.9 units of β-galactosidase activity, whereas the D133A reporter, which is defective in translation arrest *in vitro* (**Fig. 1b**), exhibited 17.9 units (**Fig. 1d**). The substantially lower β-galactosidase activity of the wild-type reporter indicates that LudP efficiently induces translation arrest in *B. subtilis* cells. In contrast, when the same reporters were expressed in *E. coli*, the difference in β-galactosidase activity between the wild-type and D133A reporters was less than two-fold (**Fig. 1e**), suggesting that LudP induces only weak translation arrest in *E. coli*. These *in vivo* observations are fully consistent with the results obtained *in vitro* using the PURE systems (**Fig. 1b, c**).

### Identification of the LudP stalling site

To determine the stalling site of LudP, we performed toeprinting analysis (**Fig. 1f**). The *gfp-ludP-lacZα* reporter was transcribed and translated using *B. subtilis* ribosomes, and reverse transcription was carried out using a primer complementary to the reporter mRNA. A ribosome stalled on the mRNA blocks reverse transcriptase, generating a complementary DNA (cDNA) fragment of a specific length. The size of the toeprint product was determined by capillary electrophoresis-based fragment analysis using Sanger sequencing products generated with the same primer as size markers. As negative controls, the same analysis was performed using reactions treated with the translation inhibitor chloramphenicol (CHL) and the arrest-defective D133A mutant. A toeprint signal of 175 nt was detected exclusively in the wild-type sample in the absence of CHL (**Fig. 1f**; cyan triangle). The size of this toeprint product indicates that the ribosome stalls with the Lys136 and Glu137 codons occupying the P and A sites, respectively (**Fig. 1f**).

### LudP-mediated translation arrest is released in a signal-dependent manner

Because LudP is encoded immediately upstream of *secDF* (**Supplementary Fig. 1b**), we hypothesized that it functions as a Sec-pathway monitor, like other arrest peptides encoded upstream of *sec* genes. A defining feature of Sec-monitoring arrest peptides is that their translation arrest is released during Sec-dependent translocation of the nascent chain^5,7^. We therefore examined whether LudP-mediated translation arrest is similarly released upon its own secretion.

To test this possibility, we constructed a translational fusion in which *lacZ* was fused to the 3′ end of an intact *ludP* coding sequence containing its N-terminal signal peptide (**Fig. 1g**). Release of translation arrest allows translation to proceed into the downstream *lacZ* sequence, resulting in increased β-galactosidase activity. Because LacZ is not expected to fold properly in the extracytoplasmic space^15^, the twelfth transmembrane segment of *E. coli* MalF was inserted between LudP and LacZ as a stop-transfer sequence to retain the LacZ moiety in the cytoplasm (**Fig. 1g**, TMD). If LudP-mediated arrest is released during secretion, the wild-type reporter should exhibit β-galactosidase activity comparable to that of the arrest-defective D133A mutant. Consistent with this prediction, the wild-type *ludP–malF*(TM12)–*lacZ* reporter produced β-galactosidase activity comparable to that of the D133A reporter in *B. subtilis* (13.8 vs 16.3 units; **Fig. 1g**). These results suggest that translation arrest by intact LudP is efficiently released in *B. subtilis* cells.

To determine whether arrest release depends on the signal peptide, we constructed a reporter lacking part of the N-terminal signal sequence (residues 6–25; ΔSS). The ΔSS reporter exhibited significantly lower β-galactosidase activity than the wild-type reporter (5.6 units) (**Fig. 1g**). In contrast, introduction of the D133A mutation into the ΔSS reporter restored high β-galactosidase activity (15.2 units). These results indicate that LudP-mediated translation arrest is released in a signal sequence-dependent manner, consistent with the behavior of other Sec-monitoring arrest peptides whose arrest is released during protein secretion^5,7^.

### Identification of the minimal ribosome stalling sequence of LudP

To define the minimal sequence required for LudP-mediated translation arrest, we performed a systematic internal frameshift mutational analysis. Internal frameshift mutations were introduced into different regions beginning at residue 98 of the *gfp-ludP(16–147)-lacZ* reporter, replacing the corresponding LudP sequences with unrelated amino acid sequences (**Fig. 2a**; red). Reporter variants in which residues 98–112, 98–117, or 98–122 were replaced exhibited β-galactosidase activities comparable to that of the wild-type reporter (**Fig. 2a**), indicating that these regions are dispensable for translation arrest. In contrast, replacement of residues 98–127 resulted in a marked increase in β-galactosidase activity, indicating loss of arrest activity. These results suggest that the C-terminal region beginning at Met123 is sufficient to mediate robust translation arrest. Because residues located downstream of the stalling site are not expected to contribute to arrest, these results further indicate that the minimal arrest element of LudP is contained within the 15-residue segment spanning Met123 to Glu137.

**Fig. 2.**
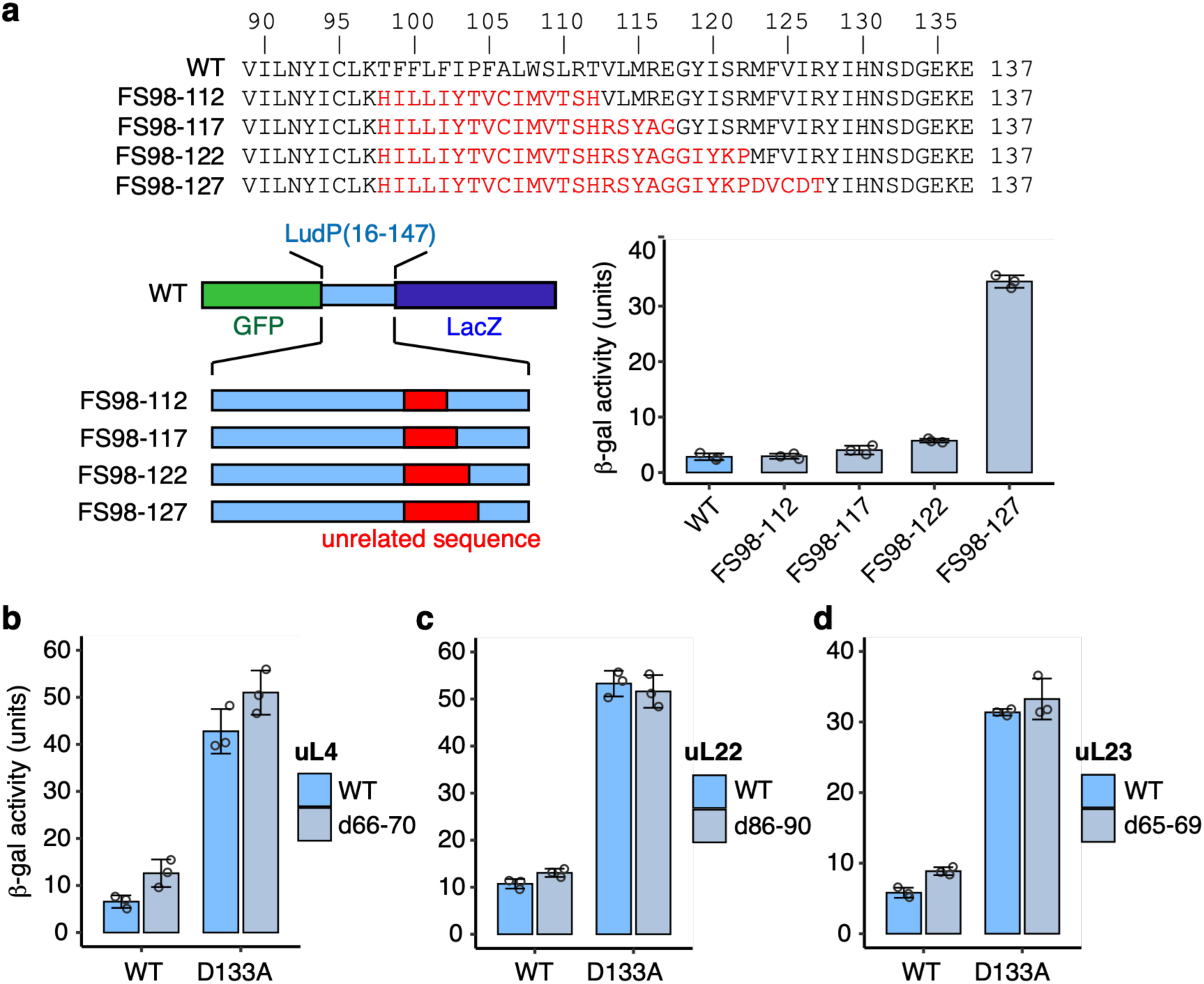
Identification of the minimal ribosome stalling sequence of LudP. **a.** Internal frameshift mutational analysis of LudP. Internal frameshift mutations were introduced into regions beginning at residue 98 of the *gfp-ludP(16–147)-lacZ* reporter. β-galactosidase activity (mean ± s.d., n = 3, biologically independent cultures) was measured in *B. subtilis* cells harboring the wild-type (WT) or mutant derivatives of the *gfp-ludP(16-147)-lacZ* reporter. The regions replaced by internal frameshift mutations and the resulting amino acid sequences are shown in red. **b-d**. β-galactosidase activity (mean ± s.d., n = 3, biologically independent cultures) of *B. subtilis* cells harboring the wild-type (WT) or D133A derivative of the *gfp-ludP(16-147)-lacZ* reporter in strains carrying or lacking loop deletions in uL4 (d66–70; b), uL22 (d86–90; c) or uL23 (d65–69; d).

### NPET mutations have little effect on LudP-mediated translation arrest

The ribosomal proteins uL4 and uL22 project into the nascent polypeptide exit tunnel (NPET) and together form its constriction site. To examine whether LudP-mediated translation arrest depends on interactions with uL4 or uL22, we performed reporter assays using *B. subtilis* strains carrying five-residue deletions in the tunnel-exposed loop regions of uL4 or uL22 (residues 66–70 and 86–90, respectively). We also examined an analogous deletion mutant of uL23 lacking residues 65–69, which is located near the exit of the NPET. Expression of the *gfp-ludP(16-147)-lacZ* reporter (**Fig. 1a**) in the uL4 mutant resulted in only a slight increase in β-galactosidase activity compared with the wild-type strain, and the activity remained much lower than that of the arrest-defective D133A mutant (**Fig. 2b**). Likewise, mutations in uL22 or uL23 had little effect on β-galactosidase activity (**Fig. 2c, d**). These results suggest that LudP-mediated translation arrest depends little, if at all, on interactions with uL4, uL22, or uL23.

### Comprehensive mutational analysis of the LudP arrest element

To systematically identify residues required for LudP-mediated translation arrest, we first performed alanine-scanning mutagenesis of the region encompassing the minimal arrest element (residues 121–137). Wild-type and alanine-substituted *gfp-ludP(16-147)-lacZ* reporter strains were cultured, and β-galactosidase activity was measured. Whereas the wild-type reporter exhibited approximately 5 units of β-galactosidase activity, alanine substitutions at I126, Y128, I129, H130, D133, G134, or K136 resulted in activities exceeding 20 units (**Fig. 3a**), indicating that alanine substitution at these positions severely impairs translation arrest. In contrast, alanine substitutions at S121, R122, M123, R127, N131, S132, or E137 produced less than 10 units of β-galactosidase activity (**Fig. 3a**), indicating that these positions largely tolerate alanine substitution.

**Fig. 3.**
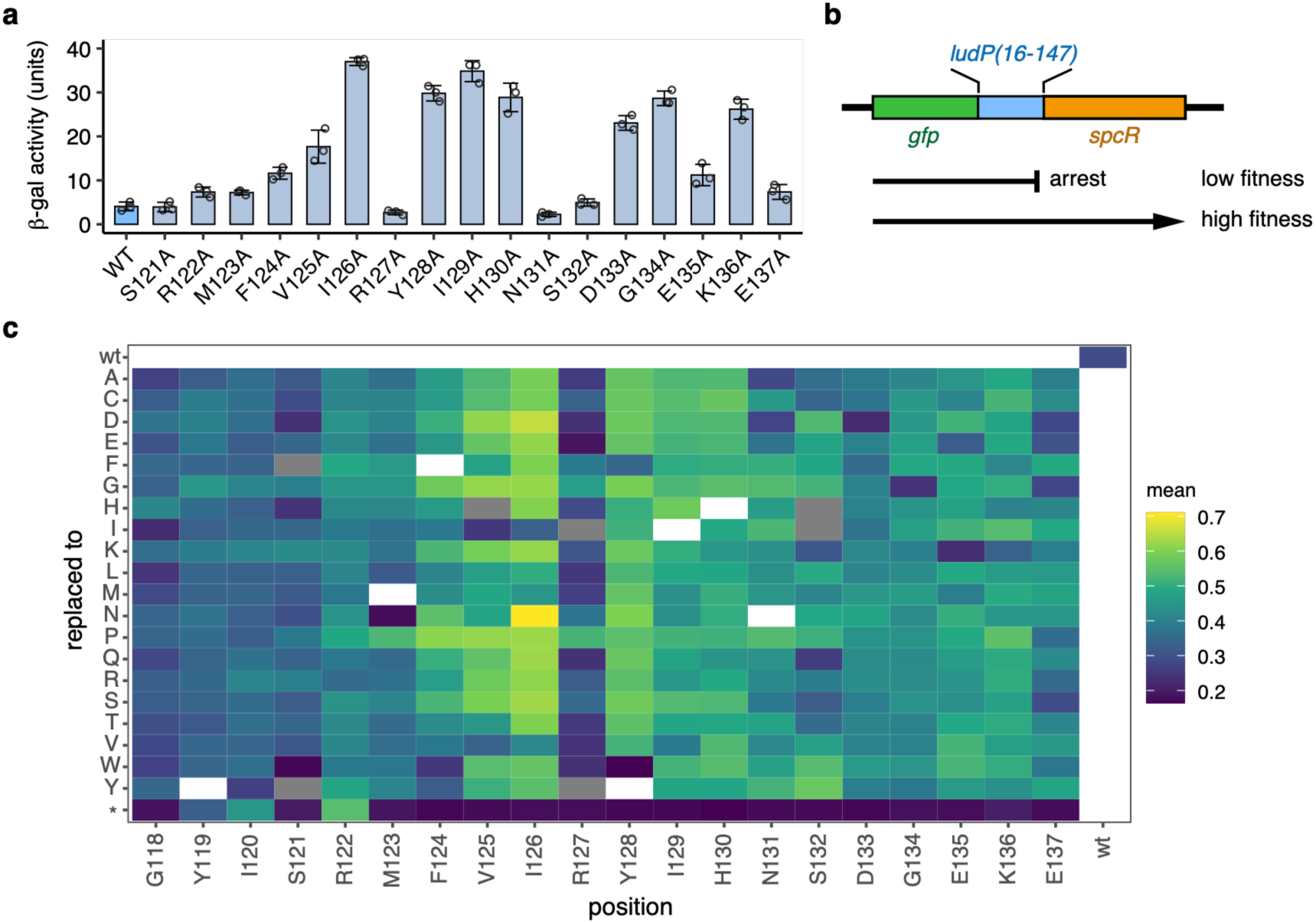
Identification of residues crucial for LudP-mediated translation arrest. **a.** Alanine-scanning mutagenesis of LudP. β-galactosidase activity (mean ± s.d., n = 3, biologically independent cultures) was measured in *B. subtilis* cells harboring the wild-type (WT) or single alanine-substitution derivatives of the *gfp-ludP(16-147)-lacZ* reporter. **b**. Schematic representation of the *gfp-ludP(16-147)-spcR* reporter used for DMS analysis shown in (c). **c.** DMS analysis of LudP. Relative fitness (mean of two biological replicates) of each mutant is shown as a heat map. Residue numbers and the corresponding wild-type amino acids are indicated at the bottom, and substituted amino acids are shown on the left. Yellow and blue indicate higher and lower fitness, respectively. White indicates wild-type. Gray indicates mutants with insufficient read counts.

Because residues that tolerated alanine substitution might nevertheless be sensitive to other amino acid substitutions, we next performed deep mutational scanning (DMS), as performed previously for the CliM arrest peptide^8^. For this purpose, we constructed a *gfp-ludP-spcR* reporter in which the spectinomycin resistance gene (*spcR*) was fused in frame to the 3′ end of *gfp–ludP(16-147)* (**Fig. 3b**). Mutations that impair translation arrest are expected to permit translation of the downstream *spcR* gene, thereby increasing cellular fitness in the presence of spectinomycin. We generated a random mixed mutant library in which each clone carried a single amino acid substitution within residues 118– 137 of LudP. The library was introduced into *B. subtilis* and cultured in the presence or absence of spectinomycin. Genomic DNA was isolated from the resulting cell populations, and the identity and abundance of each mutant were quantified by deep sequencing. Relative fitness values were calculated from these data and visualized as a heat map (**Fig. 3c** and **Supplementary Fig. 3**). Consistent with the alanine-scanning results, residues whose alanine substitutions strongly impaired translation arrest generally remained intolerant to substitution with other amino acids. In contrast, among residues that tolerated alanine substitution, most substitutions at S121, R122, M123, or R127 produced only minor effects on arrest. However, substitutions other than alanine at N131, S132, and E137 frequently impaired translation arrest, indicating that although alanine is tolerated at these positions, their side-chain properties are not completely dispensable.

### The identity of the A-site residue contributes to the stability of LudP-mediated translation arrest

In the stalled ribosome, Glu137 of LudP occupies the A-site. Our DMS analysis suggested that substitutions at this position differentially affect translation arrest. To validate this observation using a more quantitative assay, we analyzed E137 substitution mutants with the *gfp-ludP(16-149)-lacZ* reporter.

Substitution of Glu137 with Asp or Arg resulted in β-galactosidase activities comparable to that of the wild-type reporter (**Fig. 4a**). In contrast, substitution with Tyr or Phe produced β-galactosidase activities comparable to those of arrest-defective mutants (D133A and I126A). These results demonstrate that the identity of the A-site residue contributes to the stability of LudP-mediated translation arrest. This finding is also consistent with the biased amino acid distribution observed at the corresponding position among LudP homologs (**Supplementary Fig. 1c**).

**Fig. 4.**
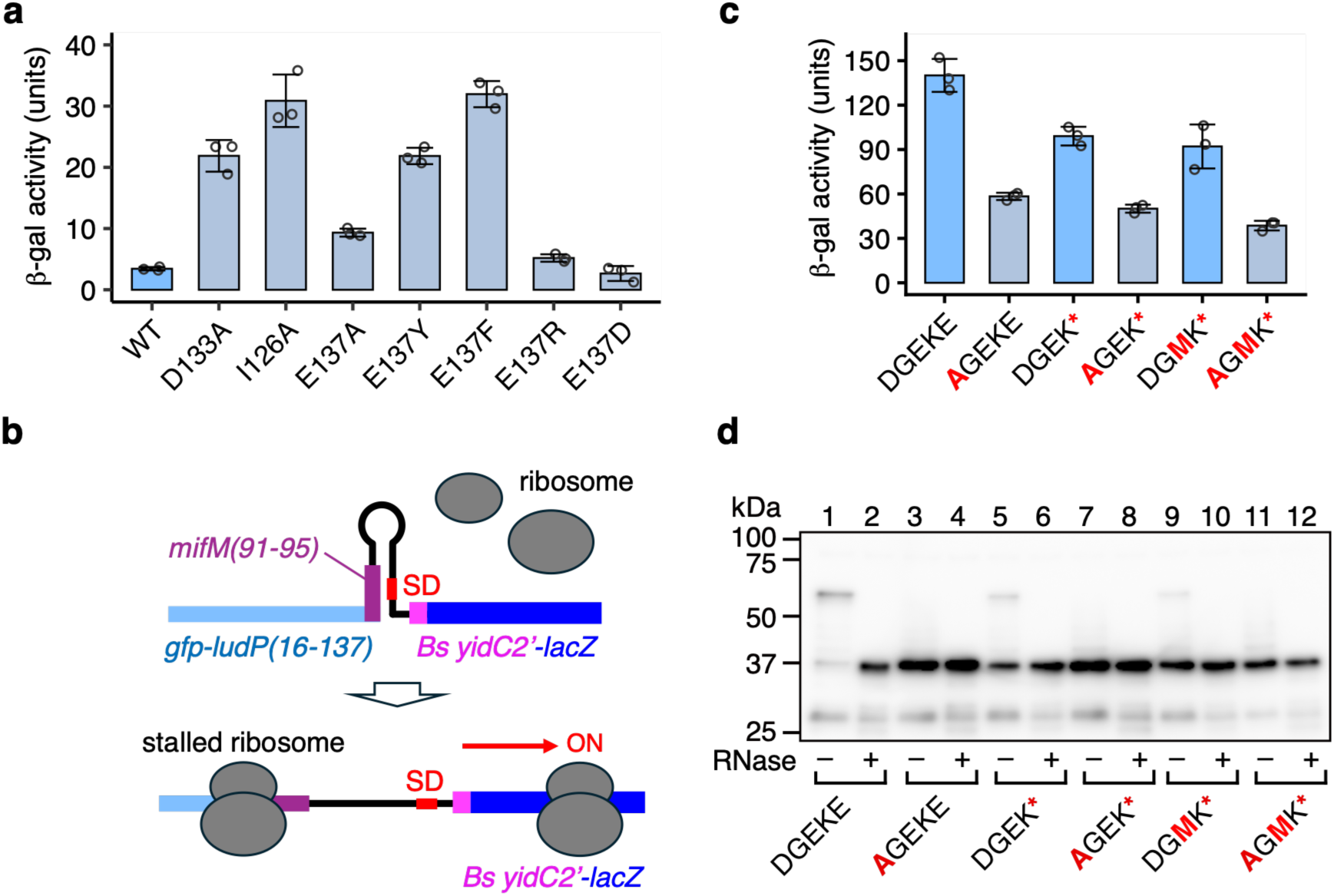
The identity of the A-site residue affects the stability of LudP-mediated translation arrest. **a**. A-site residue mutagenesis of LudP. β-galactosidase activity (mean ± s.d., n = 3, biologically independent cultures) was measured in *B. subtilis* cells harboring the wild-type (WT) or E137-substituted derivatives of the *gfp-ludP(16-147)-lacZ* reporter. **b**. Schematic representation of the *gfp-ludP-mifM-yidC2’-lacZ* reporter. **c**. β-galactosidase activity (mean ± s.d., n = 3, biologically independent cultures) was measured in *B. subtilis* cells harboring the wild-type (DGEKE), D133A (AGEKE), E137-to-stop (DGEK* and AGEK*), or DGMK-stop (DGMK* and AGMK*) variants of the *gfp-ludP-mifM-yidC2’-lacZ* reporter. **d**. *In vitro* translation assay of LudP E137-to-stop and uDF_DGMK-stop variants. Wild-type (DGEKE), D133A (AGEKE), E137-to-stop (DGEK* and AGEK*), and DGMK-stop variants (DGMK* and AGMK*) of the *gfp-ludP* construct were translated in Bs PURE at 37°C for 20 min. Translation products were analyzed by immunoblotting with an anti-GFP antibody. A portion of each sample was treated with RNase A to digest the tRNA moiety of peptidyl-tRNA.

### LudP can induce translation termination arrest

To determine whether LudP can also induce translation termination arrest, we designed a *lacZ* reporter capable of detecting both elongation and termination arrest in vivo by modifying a previously described *B. subtilis mifM–yidC2′–lacZ* reporter^6^. In this reporter, the SD sequence of *yidC2* is sequestered within a stem-loop structure formed by the *mifM-yidC2* mRNA, thereby repressing translation initiation. Translation arrest near the 3′ end of *mifM* maintains this stem-loop in an unwound conformation, exposing the SD sequence and promoting translation of the downstream *yidC2′–lacZ* reporter. We replaced the *mifM* arrest region with the coding sequence for GFP–LudP(16–137), positioning the LudP arrest sequence immediately upstream of the stem-loop (**Fig. 4b**). In this reporter, translation arrest by LudP is expected to induce expression of the downstream *B. subtilis yidC2′–lacZ* fusion by disrupting the inhibitory stem-loop. Reporter variants carrying the E137-to-stop mutation (DGEK*) or the arrest-defective D133A mutation were also constructed, introduced into *B. subtilis*, and analyzed by β-galactosidase assay (**Fig. 4c**). The wild-type reporter (DGEKE) exhibited high β-galactosidase activity (140.1 units), whereas the arrest-defective AGEKE reporter showed substantially lower activity (58.4 units; **Fig. 4c**), confirming that β-galactosidase activity reflects the efficiency of translation arrest in this reporter system. The DGEK* reporter exhibited an intermediate activity (99.0 units), significantly lower than that of the wild-type reporter but substantially higher than that of the AGEK* reporter (50.1 units). These results suggest that LudP can induce termination arrest, albeit less efficiently than elongation arrest.

To examine LudP-mediated termination arrest in vitro, we used the same reporter constructs for in vitro translation in Bs PURE (**Fig. 4d**). Whereas the wild-type construct predominantly produced the RNase-sensitive peptidyl-tRNA product, the DGEK* variant produced a reduced amount of peptidyl-tRNA together with an increased amount of released full-length product migrating near the 37-kDa marker (**Fig. 4d; lanes 1, 2, 5, and 6**). These results indicate that DGEK* induces translation termination arrest, although less efficiently than the elongation arrest observed with wild-type LudP. In contrast, the AGEK* mutant failed to produce an RNase-sensitive product (lanes 7, and 8), confirming that termination arrest depends on the LudP arrest sequence.

### Interchangeability of the LudP and uDF_DGMK-stop arrest motifs

We previously identified uDF_DGMK-stop as an arrest peptide in a subset of phylum Bacillota^12^ – the same bacterial phylum in which LudP was identified (**Supplementary Fig. 1a**). uDF_DGMK-stop contains a conserved DGMK motif at its C-terminus. Although the evolutionary relationship between LudP and uDF_DGMK-stop remains unclear, their sequence similarity raises the possibility that they share mechanistic features underlying translation arrest. To test this possibility genetically, we examined whether replacing the C-terminal region of LudP (D_133_GEKE_137_) with the DGMK* sequence would preserve arrest activity.

We examined the arrest activity of the LudP-DGMK* variant using the *mifM-yidC2’-lacZ* reporter system (**Fig. 4b**). The DGMK* reporter exhibited β-galactosidase activity that was lower than that of the wild-type reporter but comparable to that of the DGEK* reporter (**Fig. 4c**). In contrast, the AGMK* reporter showed β-galactosidase activity comparable to that of the other arrest-defective mutants. In the in vitro translation assay, the DGMK* variant accumulated both an RNase-sensitive peptidyl-tRNA species and a released full-length product in proportions comparable to those observed for the DGEK* variant (**Fig. 4d**; lanes 9 and 10). In contrast, the AGMK* variant accumulated only the released product (lanes 11 and 12). These results indicate that replacing the C-terminal region of LudP with the DGMK* sequence preserves translation arrest activity at a level comparable to that of the DGEK* variant.

We previously found that replacing the stop codon of uDF_DGMK-stop with a sense codon markedly impaired translation arrest^12^, raising the possibility that replacing the DGMK* sequence of uDF_DGMK-stop with the corresponding DGEKE sequence from LudP would likewise impair arrest. Indeed, both *in vivo* and *in vitro* analyses showed that the DGEKE variant of uDF_DGMK-stop exhibited markedly reduced arrest activity (**Supplementary Fig. 4**). Together, these results demonstrate that although the arrest motifs of LudP and uDF_DGMK-stop are partially interchangeable, each has evolved to maximize translation arrest stability within its own sequence context.

### Cryo-EM structure of a LudP-stalled ribosomal complex

To gain further insight into the molecular mechanism by which LudP induces translational arrest, we determined a structure of a LudP-stalled ribosomal complex (LudP-SRC) using single particle cryo-EM (**Fig. 5a**). Because *L. xylanolytica* LudP efficiently stalls translation on *B. subtilis*, but not *E. coli*, 70S ribosomes (**Fig. 1b-e**), we used a home-made *B. subtilis* cell-free *in vitro* translation system to generate *L. xylanolytica* LudP-SRC, as employed previously to generate *B. subtilis* MifM-, ApdA-, and CliM-SRCs^8,16,17^. The LudP-SRC was formed with full-length LudP translon (amino acids 1-147) and N-terminal FLAG-tag, which was used for purification (see **Methods**). The LudP-SRC was subsequently applied to cryo-grids and a total of 12,642 micrographs were collected on a Titan Krios transmission electron microscope equipped with a K3 direct electron detector, yielding 254,694 ribosomal particles after 2D classification (**Supplementary Fig. 5**). Following multiple rounds of 3D classification, three defined subpopulations of 70S ribosomes emerged; two major non-rotated states with tRNA in the P-site, and either a vacant A-site (61%; 155,579 particles) or A-tRNA bound in the A-site (31%, 79,551), as well as one minor rotated state bearing A/P- and P/E-site tRNAs (4.6%, 11,650 particles) (**Supplementary Fig. 5**). These three states were initially refined to average resolutions of 3.2 Å, 3.3 Å and 4.5 Å, respectively (**Supplementary Fig. 5**), however, initial inspection of the cryo-EM maps of the non-rotated P-tRNA only and rotated A/P- and P/E-tRNA states revealed a complete absence of density for any nascent polypeptide chain (NC), possibly due to flexibility and/or heterogeneity (see **Methods**), and therefore these states were not considered further. By contrast, strong density for the NC was observable within the non-rotated P- and A-tRNA containing state and therefore this state was refined further, yielding a cryo-EM map of the LudP-SRC with A- and P-tRNA, with an average resolution of 2.6 Å, and reaching 2.3 Å within the core of the large subunit (**Supplementary Fig. 6** and **Table 1**),

**Fig. 5.**
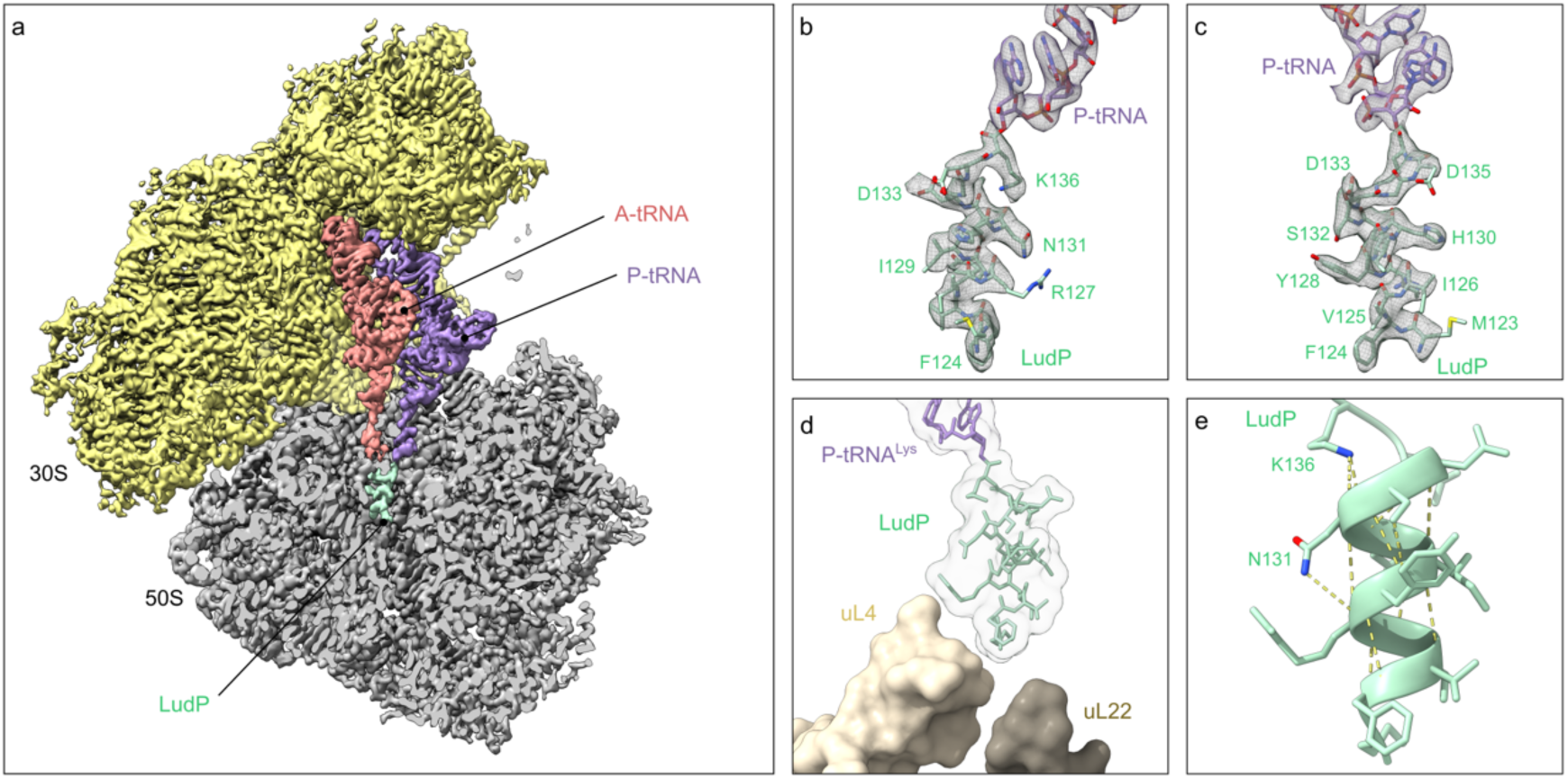
Cryo-EM structure of LudP-SRC. **a** Cryo-EM map of 3D-refined *B. subtilis* LudP-SRC with a transverse section of the large 50S subunit (grey) to reveal density for the LudP NC (mint), P-tRNA (lavender), A-tRNA (salmon) and small 30S subtunit (yellow). **b** Cryo-EM map density (black mesh) for P-site tRNA with attached LupD NC of the 3D-refined *B. subtilis* LudP-SRC. The P-tRNA (lavender) bears the LudP NC (mint). **c** Same representation as in b but turned 90° to reveal density of more LudP residues. **d** P-tRNA (lavender) and LudP (mint) in relation to uL4 (light beige), uL22 (dark beige) in surface representation. **e** Intermolecular contacts (dashed yellow lines) of LudP (mint).

**Table 1:**
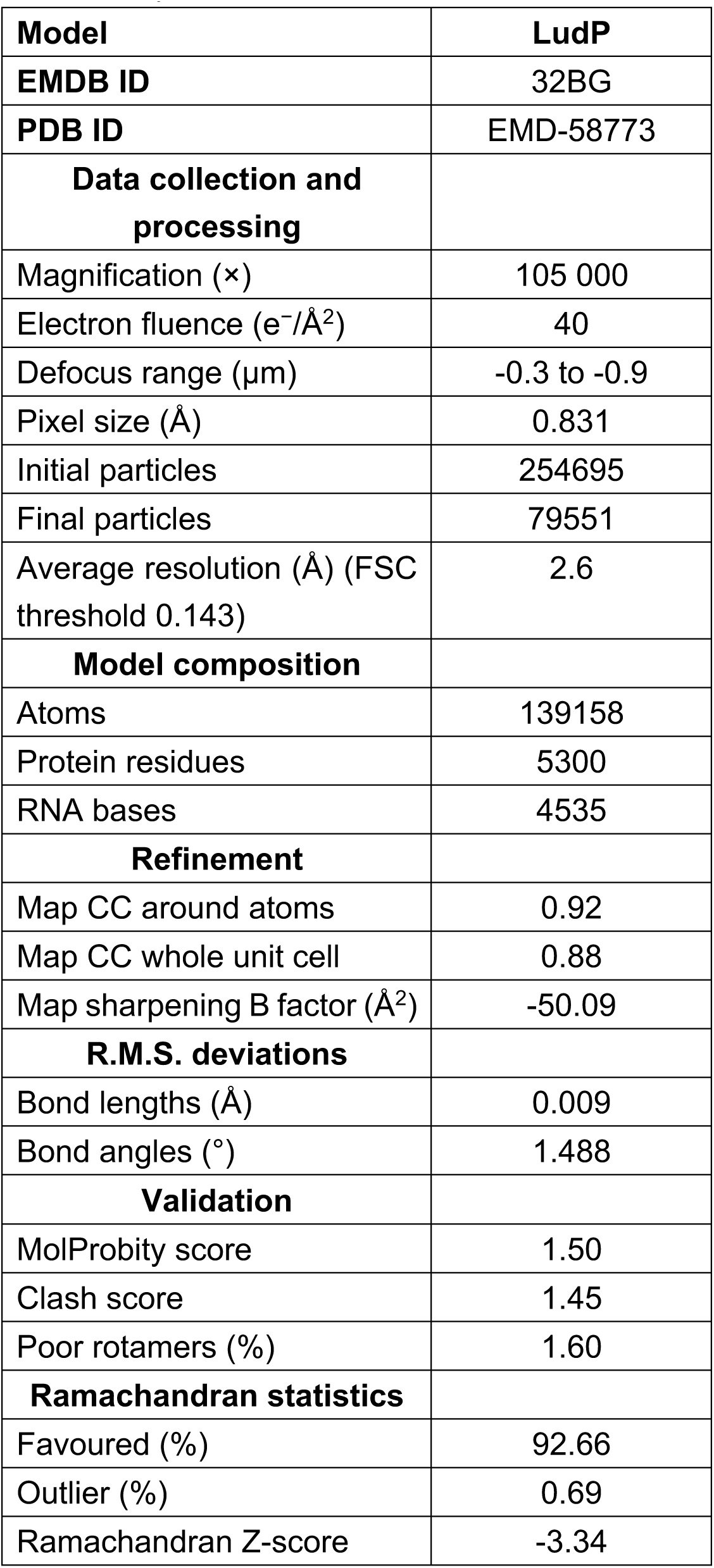
Cryo-EM data collection, refinement and validation statistics.

In the LudP-SRC with A- and P-tRNAs (**Fig. 5a**), the C-terminal region of the NC is well-resolved enabling 14 amino acids (Met123 to Lys136) to be modelled, including many side chains (exceptions were Met123, Arg127 and Asp135) (**Fig. 5b,c, Supplementary Fig. 6** and **Supplementary Movie 1**). By contrast, the density for the N-terminal region of the NC (<Met123) was poorly ordered, presumably due to flexibility, and could not be modelled. This is consistent with the biochemical analysis revealing that the C-terminal 15 residues (123-137) of LudP are sufficient to induce efficient translational arrest, whereas the N-terminal residues appear to be dispensable (**Fig. 3a**). The direct attachment of the NC via Lys136 to the P-tRNA is consistent with the toeprint analysis (**Fig. 1f**) indicating that LudP arrests translation with the Lys136 codon in the P-site. While the density for the bulk of the A-site tRNA was well-resolved and consistent with tRNA^Glu^, the CCA-end and attached Glu moiety were poorly ordered and therefore could not be modelled (**Fig. 5b,c** and **Supplementary Fig. 6**). The path of the LudP NC extends from the CCA-end of the P-tRNA, located at the PTC, into the NPET where it reaches towards the constriction formed by the tips of loops in ribosomal proteins uL4 and uL22 (**Fig. 5d**). Although the LudP NC approaches uL4 and uL22, no direct interaction was observed, consistent with biochemical assays showing that deletions within these ribosomal proteins (and uL23) do not influence the efficiency of LudP-mediated translation arrest (**Fig. 2b-d**). Careful inspection of the density and molecular model for the LudP NC revealed that residues Phe124-Asp133 adopt a standard [i + 4 → i] α-helix where each backbone nitrogen (N-H) forms a hydrogen bond with the backbone carbonyl-oxygen (C=O) of the amino acid four residues earlier (**Fig. 4d,e**). Consistently, we observe that proline substitutions within the α-helix region of LudP generally led to increased Spc resistance (**Fig. 3c**), which indicates a reduction in the stalling activities and therefore suggests that α-helix formation is important for LudP-mediated translation arrest. Overall, the ten-residue α-helix in LudP is highly reminiscent of α-helices observed at the PTC of other arrest peptides, such VemP^18^, SecM^10^, TnaC^19^, hCMV^20^, and more recently CliM^8^, although the exact length and location differs (**Supplementary Fig. 7**).

### Interaction of the LudP arrest peptide within the ribosomal tunnel

The LudP NC establishes extensive contacts with the 23S rRNA nucleotides of the NPET (**Fig. 6a-h**). This includes an intricate network of hydrogen bond interactions between the C-terminal residues of LudP and nucleotides at the PTC, namely, Lys136 with U2585 (**Fig. 6a**), Glu135 with G2061 and A2062 (**Fig. 6b**), Gly134 with U2585 and U2506 (**Fig. 6c**), and Asp133 with G2505, U2584 and C2610 (**Fig. 6d**). Many of these interactions are likely to be important since our mutagenesis analysis indicated that Asp133, Gly134 and Lys136 are critical for LudP-dependent stalling activity (**Fig. 3a,c**). We do not consider potential interactions from the side chains of Glu135 and Arg127 of LudP since both side chains are poorly resolved in the cryo-EM map, and mutations at these positions did not generally alter the arrest activity of LudP (**Fig. 3a,c**). By contrast, the side chains of His130 and Tyr128 of LudP are well-resolved and mediate potential hydrogen bond interactions with A2059 and U2609, respectively (**Fig. 6e,g**). Mutation of His130 or Tyr128 leads to strong reduction in LudP-arrest activity (**Fig. 3a,c**), suggesting that these interactions are likely to be important for stabilizing the position of the α-helix in the NPET. Additionally, the helix is stabilized in the NPET by stacking interactions between His130 of LudP, with the A2503-A2062 basepair (**Fig. 6e**), Phe124, which stacks upon the U2509-A752 basepair (**Fig. 6h**), and the side chains of Tyr128 and Ile129 of LudP, which form van der Waal and hydrophobic interactions with 23S rRNA nucleotides G2505, C2610 and U2609 (**Fig. 6f**). We note that while mutations at Phe124 are tolerated (**Fig. 3a,c**), suggesting that this stacking interaction not critical for arrest activity, mutations at Ile129 generally lead to a loss of stalling (**Fig. 3a,c**), indicating the importance of these hydrophobic interactions for translation arrest.

**Fig. 6.**
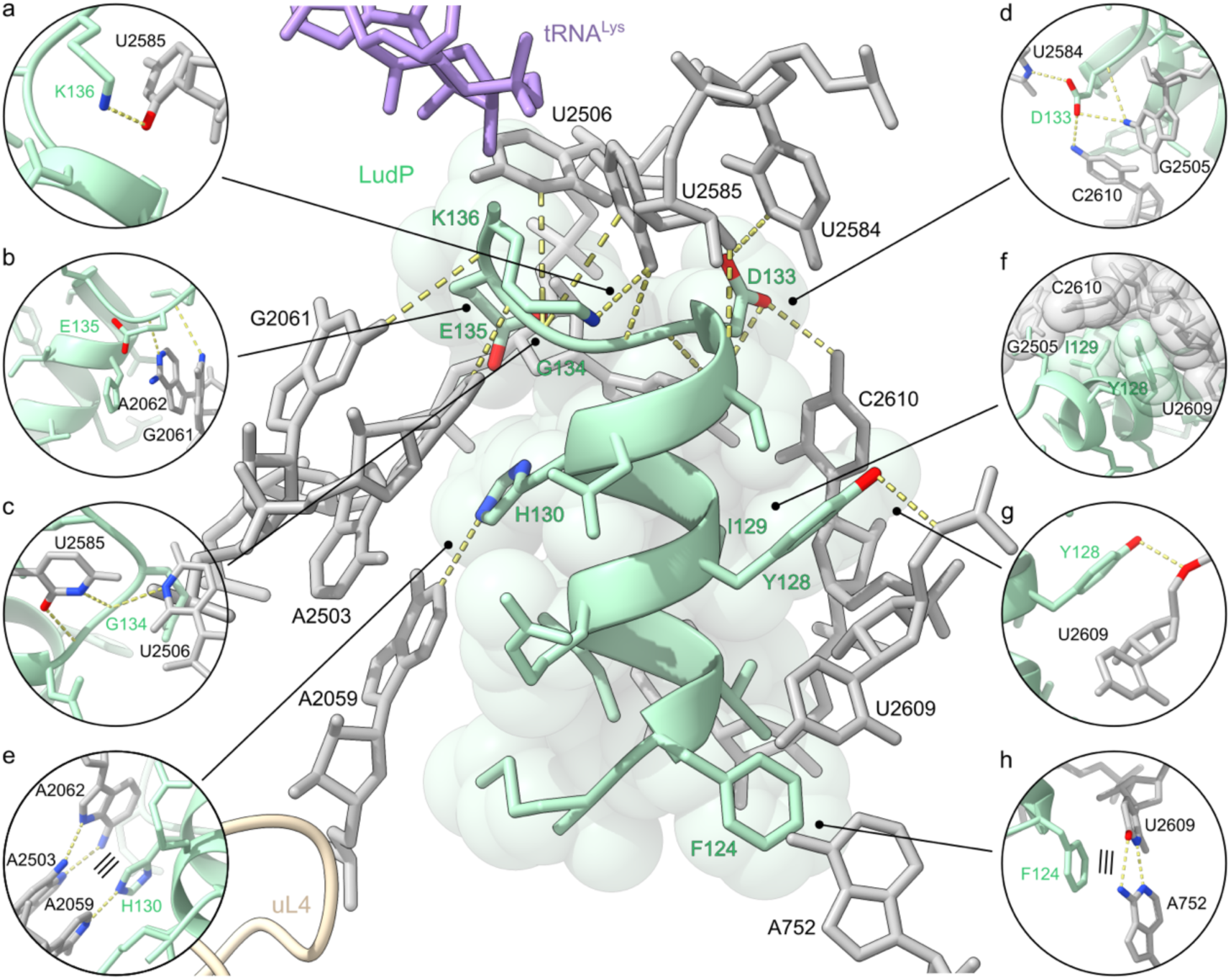
Interaction of the LudP NC with the NPET. **a** The central panel shows the LudP nascent chain (mint) attached to the P-site tRNA (lavender) and ribosomal protein uL4 (light gold). Prominent contacts with the 23S rRNA (grey) are shown in individual panels, including a hydrogen bond between the side chain of Lys136 of LudP and 23S rRNA residue U2585 (**a**), hydrogen bonds between the backbone of Glu135 and 23S rRNA residues G2061, A2062 (**b**), hydrogen bonds between the backbone of G134 of LudP and the 23S rRNA residues U2506 and U2585 (**c**), hydrogen bonds between the backbone and side chain of Asp133 of LudP and 23S rRNA residues G2505, U2585 and C2610 (**d**), a hydrogen bond between the side chain of His130 of LudP and 23S rRNA residue A2059 as well as a staking interaction between the side chain of His130 of LudP and the cooperatively interacting 23S rRNA residues A2062 and A2503 (**e**), space filling by Tyr128 and Ile129 of LudP in the cleft build by 23S rRNA residues G2505, U2609 and C2610 (**f**), a hydrogen bond between the side chain of Tyr128 of LudP and 23S rRNA residue U2609, and a stacking interaction between the side chain of F124 of LudP and the cooperatively interacting 23S rRNA residues A752 and U2609 (**h**). Contacts are shown as dashed yellow lines and stacking interactions are emphasized by three parallel lines.

### LudP induces conformational rearrangements at the PTC

The cryo-EM structure of the LudP-SRC indicates that Lys136 of LudP is attached to the P-site tRNA (**Fig. 5a-c**), suggesting that translation arrest arises because peptide bond formation cannot occur with Glu137 that is attached to the A-site tRNA. This does not appear to be due to the complete absence of binding of the Glu-tRNA to the A-site *per se* since we observe a large proportion of ribosomal particles (31%) that contain both A- and P-site tRNAs (**Supplementary Fig. 5**). However, as noted above, while the density for most of the A-site tRNA is well-resolved, the density for the CCA-end is poorly resolved, suggesting that LudP-mediated translation arrest interferes with peptide bond formation by blocking the stable accommodation of CCA-end at the PTC. To understand why the CCA-end of the A-site tRNA has not accommodated at the PTC of the LudP-SRC, we compared the conformation of the PTC of the LudP-SRC with that of the previously reported pre-attack complexes containing accommodated A- and P-site tRNAs (i.e. directly before peptide bond formation)^21^. While the general architecture of the PTC nucleotides is remarkably similar, we observe that two 23S rRNA nucleotides, namely U2585 and U2506, differ in their conformation (**Fig. 7a**). Comparing the pre-attack state (**Fig. 7b**) with the LudP-SRC (**Fig. 7c**) illustrates that the conformations of U2585 and U2506 in the pre-attack state are not compatible with the LudP-SRC because they would clash with the side chain of Asp133 of the LudP NC and therefore adopt a distinct conformation (**Fig. 7c**). Moreover, in the pre-attack state, U2506 adopts a position that is adjacent to the A-site tRNA (**Fig. 7d**), whereas the shifted position of U2506 observed in the LudP-SRC sterically clashes with the aminoacyl-moiety of the A-site tRNA (**Fig. 7e**), explaining why the CCA-end of the A-site tRNA cannot adopts its canonical position and becomes disordered. We note that the position of U2506 would also clash with the Gln side chain of the conserved GGQ motif of a release factor (RF) accommodated at the A-site^22^ (**Fig. 7f**), consistent with our biochemical analysis demonstrating that LudP also induces translation arrest when the A-site Glu codon is replaced with a stop codon (**Fig. 4**). This arrangement provides a structural explanation for the failure of productive A-site accommodation, by showing that the LudP-contact network reshapes the PTC rRNA landscape into a pre-attack-incompatible conformation.

**Fig. 7.**
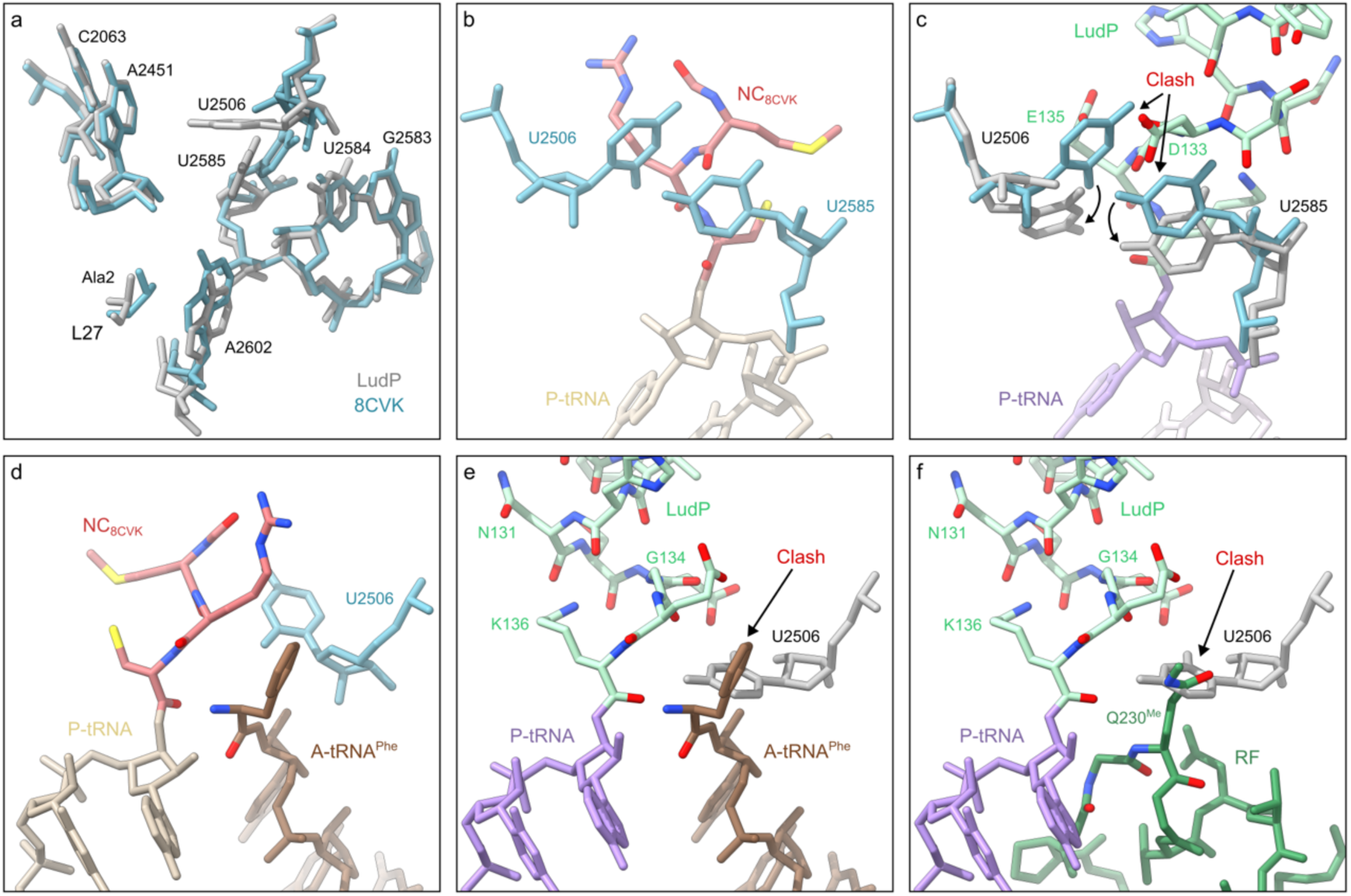
Mechanism of LudP-mediated blocking of A-site accommodation. **a** Overlay of ribosomal components involved in the proton wire required for nucleophilic attack for peptide bond formation from LudP (grey) and a pre-attack state (turquoise) (PDB ID 8CVK^21^). **b** View of the PTC of a pre-attack state (PDB ID 8CVK^21^), showing a tripeptidyl-NH-tRNA (salmon/beige) in the P-site together with PTC-forming 23S rRNA residues U2506 and U2585 (turquoise). **c** View of the PTC of LudP (mint) bound to the peptidyl-tRNA (lavender) illustrating how the nascent peptide displaces 23S rRNA residues U2506 and U2585 from their canonical positions (turquoise) to altered positions in LudP (grey). **d-f** View of the PTC of a pre-attack state (PDB ID 8CVK^21^), showing a tripeptidyl-NH-tRNA (salmon/beige) in the P-site together with PTC-forming 23S rRNA residue U2506 (turquoise) and a phenyl-NH-tRNA (brown) (PDB ID 8CVK^21^) in the A-site (**d**). The accommodation of an A-tRNA (brown) (PDB ID 8CVK^21^) (**e**) or a release factor (green) (PDB ID 9MTP^22^) (**f**) is incompatible with the displaced base 23S rRNA residue U2506 (grey) from LudP (mint). Alignments based on 23S rRNA.

### MD simulations reveal WT LudP structurally hinders U2506 pre-attack transitions

To check if, at physiological temperatures, U2506 and U2585 can reach their pre-attack conformations in the presence of WT or D133A LudP, we performed all-atom molecular dynamics (MD) simulations of a region containing all residues within 35 Å of LudP. For each variant, we carried out ten independent 2-µs simulations. WT LudP simulations were started from the LudP-SRC cryo-EM structure (containing A- and P-tRNAs). For the D133A LudP simulations, the same structure was used after in silico mutation of the Asp133 side chain to Ala. To capture the conformational landscape of the two nucleotides, we carried out a principal component analysis (PCA) of their coordinates obtained from the simulation trajectories and projected them onto the two most dominant modes of motion, i.e. principal components (**Fig. 8a,b**). In the presence of WT LudP, the U2506 motion observed in the simulations is largely orthogonal to the motion required to reach the pre-attack conformation (**Fig. 8a**, upper panel). In contrast, in the presence of D133A LudP (lower panel), we observed a motion in the direction of the pre-attack conformation, reaching very similar conformations (orange). For U2585, conformations similar to the pre-attack conformations are reached irrespective of the LudP variant. These results indicate that U2506 is hindered by WT LudP from reaching the pre-attack state while U2585 does not contribute substantially to stalling. However, the timescales of the simulations might be too short to observe a spontaneous transition of U2506 to its pre-attack conformation. Therefore, to check if LudP can adopt conformations that would not overlap with the pre-attack conformations of U2506 and U2585, we calculated the van der Waals overlap between the LudP peptide and the aligned pre-attack conformations from the trajectories. To monitor the dynamics of the peptide, we performed a PCA of the backbone atoms of the 5 C-terminal amino acids. Histograms of overlap and peptide conformational mode 1 (**Fig. 8c**) show that the D133A variant can reach lower conformational mode 1 values, corresponding to a movement of the peptide away from the PTC (black arrow, upper rightmost panel). For WT LudP, we observed frequent conformations that do not overlap with pre-attack U2585 (yellow cross) and no conformations that do not overlap with both nucleotides. In contrast, D133A LudP frequently reaches conformations that do not overlap with both pre-attack nucleotides (brown cross). In summary, our results suggest that the D133A mutation has two effects, first, the peptide can explore conformations reaching further into the tunnel than WT LudP and, second, reduction of overlap due to the smaller Ala side chain. Both effects free up the space required for U2506 to adopt the pre-attack conformation which would in turn allow full A-tRNA accommodation. This mechanism also suggests that the contribution of the helix to the stalling (**Fig. 3**) might be to prevent the peptide from moving further into the tunnel and ensuring the Asp133 position required for stalling. Together with the cryo-EM nucleotide states, the MD results support a mechanistic causality for Asp133: D133A reduces the steric/energetic constraints that otherwise prevent U2506 from adopting the pre-attack-compatible configuration.

**Fig. 8.**
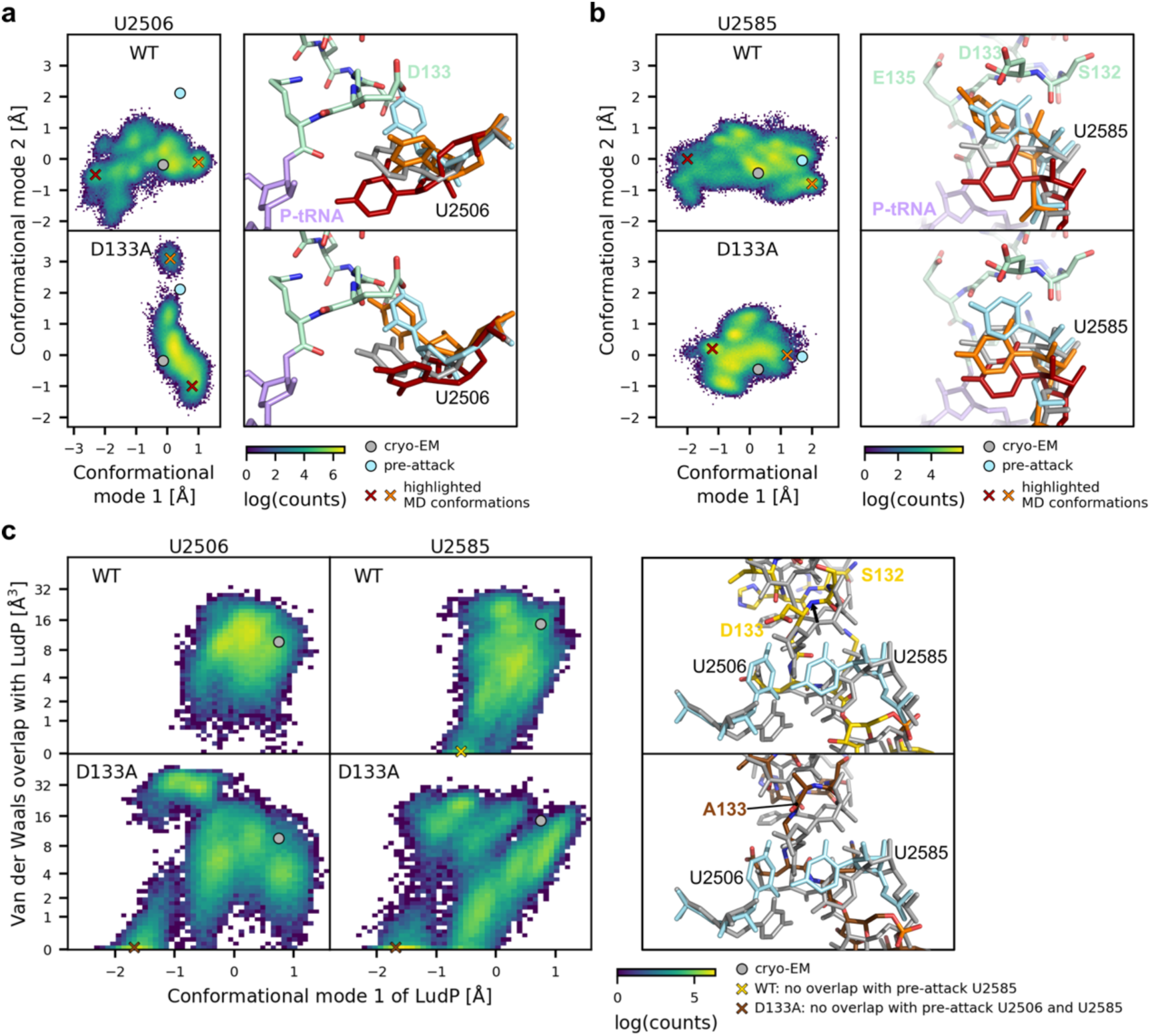
MD simulations of WT and D133A LudP. (**a**,**b**) Conformational landscape of U2506 (**a**) and U2585 (**b**) in presence of WT and D133A LudP variants obtained from MD simulations. Left, histograms of projections onto the two most dominant conformational modes with positions of cryo-EM and pre-attack conformations indicated by grey and cyan circles, respectively. Right, two nucleotide conformations that illustrate their range of motion observed in the MD simulations (red, orange; positions indicated by crosses in left panel) are shown in the context of cryo-EM conformation of P-tRNA (lavender) and LudP (mint). (**c**) Left, histograms of the dominant conformational mode of WT or D133A LudP and of the overlap volume between LudP and the pre-attack conformations of U2506 or U2585. Positions of conformations without overlap between WT LudP and pre-attack U2585 (yellow cross) and without overlap between D133A LudP and both nucleotides (brown cross). Right, the indicated conformations of WT (yellow) and D133A (brown) LudP compared to its cryo-EM conformation and pre-attack conformations of U2506 and U2585.

## Discussion

In this study, we identified LudP as an arrest peptide from a subset of bacteria belonging to the order *Lachnospirales* within the class *Clostridia*. We showed that *ludP* is encoded upstream of *secDF*, that LudP induces translation arrest near its C-terminus, and that this arrest is relieved in a signal sequence-dependent manner (**Fig. 1**). Because these characteristics are shared with previously characterized Sec-monitoring arrest peptides^5,7^, our results suggest that LudP is a member of the Sec-monitoring arrest peptide family.

Systematic mutational analyses revealed that many of the 15 amino acid residues spanning positions 123 to 137 contribute to the maintenance of translation arrest (**Fig. 2, 3**). Furthermore, replacement of many residues within this region with proline impaired arrest activity, suggesting that LudP adopts an α-helical structure in this region and that this structure plays an important role in translation arrest. Consistent with this idea, our structural analysis showed that LudP adopts a distinctive α-helical conformation within the NPET (**Fig. 5**), and that multiple residues within this helix interact with ribosomal RNA (**Fig. 6**). Formation of an α-helix within the NPET positions numerous amino acid side chains on the outer surface of the helix, thereby maximizing nascent chain-ribosome interactions and stabilizing specific nascent chain conformations within the NPET. This may explain why many arrest peptides adopt α-helical structures within the NPET^8,10,18–20,23^.

Translation arrest frequently involves proline residues located near the PTC. For example, SecM, ApcA, ApdA, ApdP, and their homologs contain proline-containing motifs, such as RAPP, RGPP, RAPG, or RAGP, near their arrest sites, in which the proline residue plays a critical role^10,16,24^. In contrast, several arrest peptides, including MifM, VemP, CliM, and LudP, do not contain a proline residue near the PTC^8,17,18^. Whereas the C-terminal regions of SecM, ApdA, and ApdP adopt essentially the same configuration around the PTC^10,16^, MifM, VemP, CliM, and LudP each assume distinct conformations^8,17,18^. These observations illustrate the remarkable diversity of translation arrest mechanisms that has evolved in bacteria.

Some proline-dependent arrest peptides, such as ApdA and ApdP, are largely unaffected by mutations in the ribosomal constriction site^16^. In contrast, the arrest activities of most proline-independent bacterial arrest peptides are reduced by mutations in uL4 or uL22^6–8,17^. Nevertheless, LudP-mediated arrest was largely unaffected by mutations in either uL4 or uL22 (**Fig. 2**). Consistent with this observation, the 15 amino acid residues required for LudP-mediated arrest are located between the PTC and the NPET constriction site, and none of these residues directly interacts with uL4 or uL22 (**Fig. 5**). In this respect, LudP represents a unique variation of the proline-independent translation arrest mechanism.

Our genetic, biochemical, structural, and computational analyses converge on a single mechanism for LudP-mediated translation arrest (**Fig. 9**). The C-terminal LudP arrest element positions Asp133 within the ribosomal peptidyl-transferase center (PTC), where it induces conformational rearrangements of 23S rRNA nucleotides, most notably U2506. In the LudP-stalled ribosome, the shifted position of U2506 sterically and energetically disfavors productive accommodation of the A-site aminoacyl-tRNA, as well as a termination release factor, if the A-site codon is mutated to a stop codon (**Fig. 9a,b**). Consistent with this model, D133A abolishes the nucleotide rearrangements and restores access to a pre-attack-compatible U2506 conformation in molecular dynamics simulations, thereby re-enabling stable A-site accommodation (**Fig. 9c**). Finally, the identity of the A-site residue is encoded by the LudP arrest element itself: E137 in the A-site promotes stable arrest, whereas substitutions at this position impair the capacity of LudP to maintain the non-productive ribosomal state. Blocking aminoacyl-tRNA or RF accommodation is a mechanism shared with MifM, VemP, and CliM^8,17,18^. However, the molecular details differ among these arrest peptides. In MifM and VemP, as in LudP, specific rRNA nucleotides act as steric barriers to A-site accommodation. In contrast, in CliM, side chains of the nascent peptide directly block the accommodation of RF or aminoacyl-tRNA. Among the arrest peptides characterized to date, the mechanism of LudP most closely resembles that of VemP^18^. Both arrest peptides form an α-helical structure within the NPET, and reposition specific rRNA nucleotides near the PTC to induce structural rearrangements in the A-site.

**Fig. 9.**
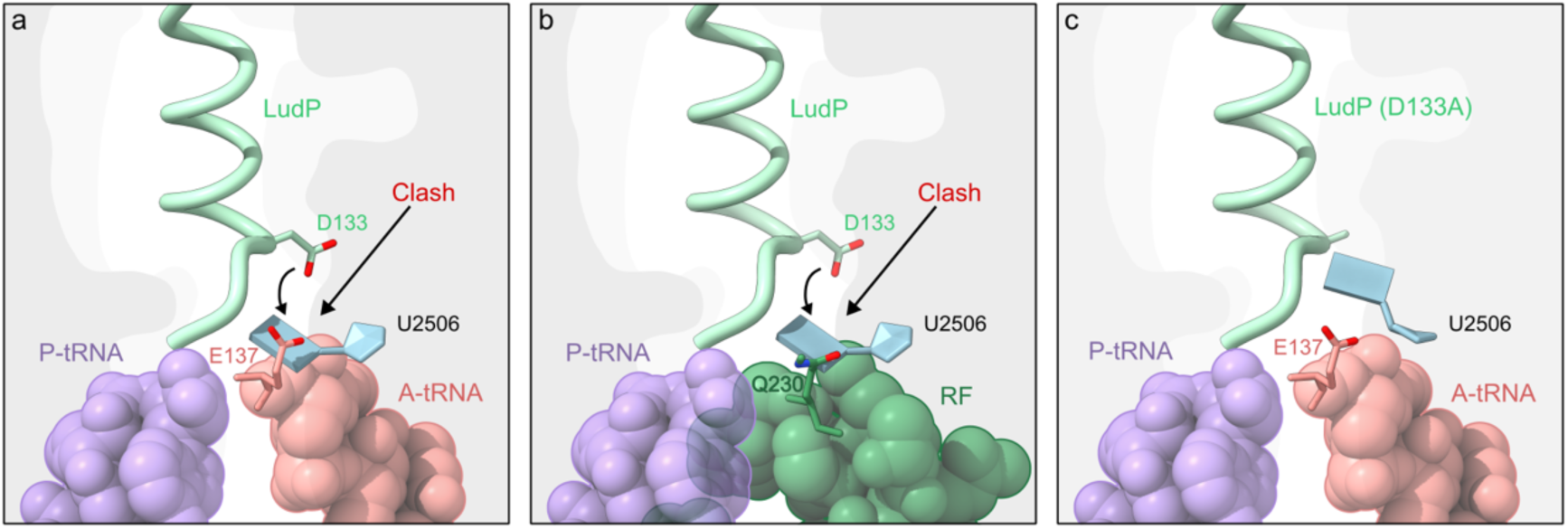
Model for LudP-mediated translational stalling. (**a-b**) LudP adopts a stable conformation in the NPET and the side chain of D133 displaces U2506 into the A-site of the ribosome. Therefore, U2506 interferes with translation elongation (a) as well as termination (b) due to a steric clash with the amino acid moiety (a) or Q230 of the RF, respectively. (**c**) D133A mutation in LudP enables transition of U2506 to its canonical pre-attack state allowing A-site accommodation in the PTC.

Although LudP blocks aminoacyl-tRNA accommodation in the A-site, efficient translation arrest is not induced with all aminoacyl-tRNAs recruited to the A-site (**Fig. 4**). Likewise, replacement of the A-site codon with a stop codon still allowed translation arrest to occur, but with reduced efficiency. Such A-site residue selectivity was not observed for CliM^8^, whereas certain substitutions at the A-site residue of VemP also reduce arrest activity^25^. However, the identities of the permissive A-site residues appear to differ between LudP and VemP. Consistent with this difference, glycine or glutamate is frequently conserved at the A-site position among LudP homologs, whereas phenylalanine or tyrosine is preferentially conserved at the corresponding position in VemP homologs. A-site residue selectivity has also been observed in several other arrest peptides. For example, the A-site proline of SecM, ApdA, and ApdP plays an active role in their arrest mechanisms, making the presence of proline at this position indispensable^10,16^. Likewise, during drug-dependent stalling of ErmDL, electrostatic repulsion between the basic residue immediately preceding the arrest site and the basic residue occupying the A-site is essential for translation arrest^26^, thereby requiring a basic amino acid at the A-site position. The mechanism underlying A-site residue selectivity in LudP and VemP remains unknown. One possibility is that, in the LudP-RNC, interference of U2506 with A-site accommodation is not completely static. Depending on the identity of the incoming aminoacyl-tRNA, accommodation may overcome the steric interference imposed by U2506, thereby reducing the efficiency of translation arrest.

To date, four members of Sec/YidC-associated arrest peptides have been identified in the bacterial phylum Bacillota. Among them, MifM and uDF_DGMK-stop are found predominantly in bacteria belonging to the class *Bacilli*, whereas CliM and LudP are found in bacteria belonging to the class *Clostridia* (**Supplementary Fig. 1a**). Thus, uDF_DGMK-stop and LudP were identified from phylogenetically distinct bacterial classes. In addition, our clustering analysis assigned them to different sequence clusters^12^. For these reasons, we have treated them as distinct arrest peptide families. Nevertheless, we do not exclude the possibility that they share a common evolutionary origin. Their partial sequence similarity and partial interchangeability suggest that they employ related molecular mechanisms. On the other hand, replacing the A-site stop codon of uDF_DGMK-stop with the glutamate codon, which is frequently found at the A-site position of LudP homologs, markedly impaired translation arrest^12^, suggesting that each has evolved to optimize translation arrest within its own sequence context.

Sec/YidC-associated arrest peptides provide an excellent example of functional convergent evolution of arrest peptides. The identification of LudP further expands this repertoire and reinforces the emerging view that highly diverse arrest peptide sequences can independently evolve to perform similar physiological functions.

## Methods

### Bacterial strains and plasmids

The bacterial strains, plasmids, and DNA oligonucleotides used in this study are listed in **Supplementary Tables 1-3**, respectively. Plasmids were constructed via standard cloning methods, including PCR using PrimeSTAR GXL (Takara) and Gibson assembly^27^. Sera-Mag Carboxylate-Modified Magnetic Particles (Cytiva, 65152105050250) was used to purify double-stranded DNA and sequencing products. The *B. subtilis* strains were constructed by transformation involving double homologous recombination between chromosomal DNA and the plasmids introduced into *B. subtilis* competent cells. The resulting recombinant clones were validated based on their antibiotic-resistance markers.

### Culture media and growth conditions

*B. subtilis* cells were cultured in LB medium. *E. coli* cells were cultured in LB medium supplemented with 100 μg/mL ampicillin. Cells were cultured at 37°C and collected for Western blotting or β-galactosidase activity assay when they reached an optical density of 0.5–1.0 at 600 nm (OD_600_).

### Purification of *B. subtilis* ribosomes

*B. subtilis* strain PY79 were grown at 37 °C in LB medium until mid-log phase. Cells were then harvested by centrifugation, washed twice with buffer I-H (10 mM HEPES-KOH pH 7.6, 15 mM Mg(OAc)_2_, 1 M KCl, 5 mM EDTA, 10 mM 2-mercaptoethanol), then once with buffer II-H (Buffer I-H with 50 mM KCl) and stored as a pellet at –80 °C. They were thawed in suspension buffer (10 mM HEPES-KOH pH 7.6, 50 mM KCl, 10 mM Mg(OAc)_2_, 7 mM 2-mercaptoethanol) with protease inhibitors (1 μg/mL PefaBlock,10 μg/mL Leupeptin, 0.7 μg/mL Pepstatin) and disrupted by passing through a microfluidizer LV1 (Microfluidics) at 16,000 psi. Cell lysate was mixed with an equal volume of suspension buffer with 3 M ammonium sulfate and kept on ice for 30 min. Cell debris and protein precipitates were removed by centrifugation (4 °C, 10,000 g, for 30 min). The supernatant was subjected to HiTrap Butyl FF column of 10 mL volume (GE) equilibrated with buffer A (20 mM HEPES-KOH pH 7.6, 1.5 M (NH_4_)_2_SO_4_, 10 mM Mg(OAc)_2_, 7 mM 2-mercaptoethanol). The column was then washed with 40 mL of buffer B (20 mM HEPES-KOH pH 7.6, 10 mM Mg(OAc)_2_, 7 mM 2-mercaptoethanol) containing 1.2 M ammonium sulfate and proteins were eluted with a 1.2 M to 0 M gradient of ammonium sulfate in buffer B. Fractions containing the ribosome were combined and loaded on the top of an equal volume of 30% sucrose cushion buffer (20 mM HEPES-KOH pH 7.6, 10 mM Mg(OAc)_2_, 30 mM NH_4_Cl, 30% sucrose, 7 mM 2-mercaptoethanol) and ultracentrifuged (4 °C, 100,000 g for 16 h) to sediment the ribosomes. The pellet was suspended with ribosome buffer (20 mM HEPES-KOH pH 7.6, 30 mM KCl, 6 mM Mg(OAc)_2_, 7 mM 2-mercaptoethanol) and stored at –80 °C until use for in vitro translation assays.

### In vitro translation and Western blotting

Bacterial reconstituted transcription–translation coupling systems^13,14^ were used in the in vitro translation assay. For in vitro translation using *E. coli* ribosomes, we utilized PUREfrex version 1.0 (GeneFrontier). For Bs PURE, purified *B. subtilis* ribosomes were used at the final concentration of 1 μM in the PURE system, without adding *E. coli* ribosomes. 2.5 U/mL of T7 RNA polymerase (Takara) was added to ensure transcription. The in vitro translation reaction was primed using the DNA templates prepared by consecutive PCR as follows. The first PCR products were amplified from respective plasmids using primers PT7-RBSkf-GFP (TAACTTTAAGAAGGAGGGAGATATACCAATGACAATGTTTGTGGGATC) and lacZ60-TAATAA-21-rv (TGGTGCCGGAAACCAGGCAAATTATTAGCGCCATTCGCCATTCAGGCT), and then used as a template for the second PCR with primers Universal primer-77-PURE (GAAATTAATACGACTCACTATAGGGAGACCACAACGGTTTCCCTCTAGAAATAATTTTGTTTAACTTTAAGAAGGAG) and lacZ60-TAATAA-21-rv. The translation reaction was carried out for 20 min at 37°C and was stopped by adding 2× SDS–PAGE loading buffer, for Western blotting. A portion of the sample was further treated with 0.2 mg/mL RNase A (Promega) at 37°C for 10 min, to degrade the tRNA moiety of the peptidyl-tRNA. The translation products were separated on a 10% polyacrylamide gel that was prepared using WIDE RANGE Gel buffer (Nacalai Tesque), according to the manufacturer’s instructions, then transferred onto a PVDF membrane (Merck, IPVH00010) and subjected to immuno-detection using antibodies against GFP (mFX75; Wako, 012-22541) or LacZ (ref: Fujiwara 2023). Images were acquired and analyzed using an Amersham Imager 600 luminoimager (GE Healthcare), and the band intensity was quantified using ImageQuant TL (GE Healthcare).

### Toeprinting assay

In vitro translation was carried out using the Bs PURE at 37°C for 20 min in the presence or absence of 0.1 mg/mL chloramphenicol. The translation reaction mixture was then mixed with the same volume of the reverse transcription mixture containing 50 mM HEPES-KOH, pH7.6, 100 mM potassium glutamate, 2 mM spermidine, 13 mM magnesium acetate, 1 mM DTT, 2 mM of oligonucleotide labeled with 6-carboxyfluorescein (6-FAM) at the 5’ end (5’–AGGGCGATCGGTGCGGGCCTCTTC– 3’, Invitrogen), 50 mM each dNTP, and 10 U/mL ReverTra Ace (Toyobo), then incubated further at 37°C for 15 min. The reaction mixture was diluted 5-fold with the NTC buffer (Macherey-Nagel), and the reverse transcription products were purified using a NucleoSpin Gel and PCR Clean-up kit (Macherey-Nagel). The reverse transcription products were eluted with 30 μL of HiDi formamide (Thermo Fisher). Samples were then mixed with 10 μL of 10-fold-diluted GeneScan 500 LIZ dye size standard (Thermo Fisher, 4322682), then heated at 96°C for 3 min just before capillary electrophoresis. The dideoxy DNA samples used as size markers for sequencing were prepared using a Thermo Sequenase Dye Primer Manual Cycle Sequencing Kit (Thermo, 79260), Thermo Sequenase Cycle Sequencing Kit (Thermo, 785001KT), or Thermo Sequenase DNA Polymerase (Cytiva, E79000Y), according to the manufacturer’s instruction, with some modifications. The DNA polymerase reaction was carried out using the same sets of template DNA and primer used for the toeprint assay. Each reaction mixture contained 0.44 μM of the 6-FAM-labeled primer, 60 μM each deoxynucleotide triphosphate (dATP, dCTP, dGTP, and dUTP), and 0.6 μM dideoxynucleotide triphosphate (either ddATP, ddCTP, ddGTP, or ddUTP). The sequencing products were purified using Sera-Mag speed beads and eluted with HiDi formamide. Next, 2 μL of a 10-fold-diluted GeneScan 500 LIZ dye size standard was added. If needed, the toeprinting product was further diluted before electrophoresis using HiDi formamide. The toeprinting and dideoxy sequencing products were then subjected to fragment analysis on a Seqstudio genetic analyzer (Thermo Fisher). Fragment data were analyzed using the GeneMapper software version 6 (Applied Biosystems), and processed further using Adobe Illustrator. The signals obtained from dideoxy sequencing were colored green (A), blue (C), black (G), and red (T), and then superposed for presentation.

### In vivo β-galactosidase assay

A 100-μL aliquot of the culture was transferred to a well in a 96-well plate, and OD_600_ was recorded. We mixed the culture with 50 μL of Y-PER reagent (Thermo Fisher) for 20 min at room temperature, to disrupt the cells. Subsequently, 30 μL of *o*-nitrophenyl-β-D-galactopyranoside (ONPG) in Z-buffer (60 mM Na_2_HPO_4_, 40 mM NaH_2_PO_4_, 10 mM KCl, 1 mM MgSO_4_, and 38 mM 2-mercaptoethanol) was added to the cell lysate, and the OD_420_ and OD_550_ were measured at 28°C every 5 min over a period of 60 min, or every two in over a period of 20 min. Arbitrary units of β-galactosidase activity were calculated using the following formula: [(1000 × V_420_ − 1.3 × V_550_) / OD_600_], where V_420_ and V_550_ are the first-order rate constants, OD_420_/min and OD_550_/min, respectively.

### Deep mutational scanning of LudP

A SpcR-reporter plasmid (*gfp-Lx_ludP(16-147)-flag-spcR*: pCH2610) was constructed by fusing a gene fragment encoding residues 16-147 of *Lx* LudP in-frame with *gfp* and *flag-spcR*. Using degenerate primers (**Supplementary Table 4**), we introduced comprehensive single-residue substitutions at positions 118-137. The resulting plasmid library was integrated into the chromosome of *B. subtilis*, pooled, and stored as glycerol stocks at -80 °C.

Aliquots of the library were inoculated into LB medium without spectinomycin (Spc) and cultured to mid-log phase (OD_600_ = 0.5, defined as t0). Cells were then subcultured into LB medium with or without Spc (250 or 500 μg/mL) and grown again to mid-log phase (t1). At both t0 and t1, cells were collected for genomic DNA extraction and a portion of the cultures was serially diluted and plated on LB agar plates to determine colony-forming units (CFU). Genomic DNA was extracted as follows. Cells were resuspended in 200 μL buffer G3 (NucleoBond Buffer Set III, Takara, Cat. No. 740603), supplemented with 4 μL of lysozyme solution (100 mg/mL; Sigma Cat. No. L6876) and 5 μL of NucleoBond Proteinase K (provided in the kit), and incubated at 37 °C for 20 min. Subsequently, 80 μL buffer G4 was added, vortexed, and incubated at 50 °C for 30 min. To this lysate, 200 μL SeraPURE beads were added to capture DNA. After two washes with 70% ethanol, DNA was eluted with 200 μL MilliQ water.

For NGS library preparation, DNA was PCR-amplified using primers listed in **Supplementary Table 5**. The PCR mixture contained 2 μL chromosomal DNA (5 ng/µL) in 40 μL GXL Fast reaction mixture (Takara). PCR was carried out for 24 cycles under the following conditions: 98 °C for 10 sec, 55 °C for 5 sec, and 68 °C for 20 sec. PCR products were purified with SeraPURE beads and eluted in 10 µL EB buffer. Purified products were separated on a 10% polyacrylamide gel using 1×TBE buffer, and the PCR bands were excised from the gel. The gel slices were homogenized with a pestle in 300 µL DNA elution buffer (10 mM Tris-HCl, pH 8.0, 300 mM NaCl, 1 mM EDTA) and vortexed overnight at 37 °C at 15,000 rpm to extract DNA. The homogenates were applied to an EconoSpin column (Gene Design) and centrifuged at 11,000 rpm for 3 min at 4 °C to remove gel debris, while recovering DNA in the flow-through. The flow-through was mixed with 600 μL NTC buffer and applied to a NucleoSpin Gel and PCR Cleanup column (Macherey-Nagel). After two washes with NT3 buffer, DNA was eluted with 17 µL NE buffer. NGS was outsourced to Rhelixa (Tokyo, Japan) and performed using the Illumina NovaSeq X Plus platform.

## DMS data analysis

The DNA libraries for Illumina sequencing were constructed with the following structure: 5’-*AATGATACGGCGACCACCGAGATCTACACAGCGCTAGACACTCTTTCCCTACACG ACGCTCTTCCGATCT*NNNNNNGGTTTTCTTAATATTGTAAGCCGCTCAAATCTA AAAAGCGTAATCCTTAATTATATATGTTTAAAAACATTCTTCTTATTTATACCGT TTGCATTATGGTCACTTCGCACCGTTCTTATGCGGGAG<u>GGATATATAAGCCGGATGTTTGTGATACGTTATATTCATAATTCCGACGGAGAAAAAGAA</u>GCAGCAGG CAGGCTATGGCAGATTGAAAATGACTATAAAGACGACGACGACAAAGGTNN NNNN*AGATCGGAAGAGCACACGTCTGAACTCCAGTCACCCGCGGTTATCTCGTAT GCCGTCTTCTGCTTG*-3’. Both terminal regions indicated in italics correspond to Illumina adapters: the 5′ end contains the P5 adapter, the i5 index, and the binding site for Read 1 sequencing primer, while the 3′ end contains the P7 adapter, the i7 index, and the binding site for Read 2 sequencing primer. Each “NNNNNN” represents a six-nucleotide randomized sequence. Within these regions, the trinucleotide GGT (designated bc5) and the trinucleotide GGT (designated bc7) indicated in bold serve as barcodes. The intervening sequence between bc5 and bc7 constitutes the reporter sequence, and the underlined sequence correspond to codons 118-137 of wild type *ludP*, in which each codon was replaced with NNK random codon. Sequencing was carried out with 150-bp paired-end reads. The forward and reverse reads overlapped by 45 bp, allowing precise determination of the *ludP* region. Raw sequencing reads were demultiplexed according to their barcode sequences using Cutadapt (version 5.2) ^28^. Demultiplexed paired-end reads were subsequently merged using fastp (version 0.23.4)^29^. Bases with Phred quality scores below 20 were considered low quality, and reads containing more than 30% low-quality bases, an average Phred quality score below 20, or more than five ambiguous (N) bases were discarded. Reads were then assigned to each mutated sequence variant and counted using R (version 4.3.3) with the packages ShortRead (version 1.60.0)^30^ and tidyverse (version 2.0.0)^31^. The growth rate of each variant was calculated using the following equation: growth rate = log_2_[(CFU_t1_ / CFU_t0_) × (RPM_t1_ / RPM_t0_)] / culture hour, where *CFU*represents colony forming units, *RPM* represents reads per million, and *Culture hour* indicates the incubation time between *t0* and *t1*. The relative fitness under 500 μg/mL spectinomycin (Spec) was calculated by dividing the growth rate under the antibiotic condition by that under the no-antibiotic condition. The experiments were performed independently twice, and the mean relative fitness values were plotted as a heatmap using only variants with at least eight reads. The reproducibility between biological replicates is shown in **Supplementary Fig. 3**.

### Phylogenetic analysis

A cluster containing 22 LudP-related amino acid sequences was selected from candidate arrest peptide clusters identified in our previous study^12^. These sequences were aligned using MAFFT v7.526, and a profile hidden Markov model (HMM) was constructed from the alignment using hmmbuild (HMMER v3.4). The profile was used to search a local protein database containing sequences from 96,821 GTDB release 232 representative bacterial genomes using hmmsearch (HMMER v3.4), with a full-sequence E-value threshold of 0.01 and a conditional domain E-value reporting threshold of 0.01. Sequences retrieved in the first search were combined with the original 22 sequences, and identical amino acid sequences were removed, yielding 75 sequences. These sequences were aligned using MAFFT, and a new profile HMM was constructed for a second search against the same protein database using the same thresholds. Hits from the two search rounds were combined, and duplicate protein IDs were removed. The two datasets were then merged, and entries with identical full-length amino acid sequences were collapsed into a single representative. Genomic context was retrieved using the protein ID lists before sequence-based deduplication. The genomic positions and orientations of neighboring genes were obtained from the corresponding GFF annotation files. The nonredundant LudP-related sequences and the proteins encoded by their downstream genes were clustered separately using MMseqs2 v18.8cc5c easy-cluster with the parameters --min-seq-id 0.3, -c 0.7, and --cov-mode 0. Downstream protein clusters were assigned functional annotations based on existing protein annotations. LudP-related sequences whose downstream gene products belonged to a cluster annotated as SecDF were retained as putative LudP homologs. The selected LudP homologs were aligned using MAFFT. A sequence logo of the alignment region corresponding to residues 100– 140 of the reference LudP sequence (WP_104436629.1) was generated using the R package ggseqlogo v0.2.2. The distribution of putative LudP homologs was mapped onto the GTDB release 232 reference phylogeny for the phylum Bacillota, which was visualized and annotated using the R package ggtree v3.16.3.

### PCR and in vitro transcription

PCR reaction (Q5 High-Fidelity DNA Polymerase (NEB), Q5 Reaction buffer (NEB)) was used with primers fwd_T7_SD_FLAG_Lxyl (5’-GCGAATTAATACGACTCACTATAGGGAGAAAAGGAGGAAAACATATGGACTA CAAAGACGACGACGACAAATTGGAAACCAGGAGGAAAAAATCTGG-3’) and rev_4xSTOP_Lxyl (5’-TTATTATTATTAATTTTCAATCTGCCATAGCCTGCC-3’) on the vector carrying LudP ORF to generate the amplified DNA sequence (5’-GCGAAT<u>TAATACGACTCACTATAGG</u>GAGA<u>AAAGGAGG</u>AAAACAT<u>ATGGACTA</u> <u>CAAAGACGACGACGACAAA</u>TTGGAAACCAGGAGGAAAAAATCTGGGATACA TCGTCTGATACTGGCATGTTTTTTTTGTACGGCAGTATTAATGACACAACTGAATACGGTTGATGTGGTCAAGAGCCTGGAACATCCTGGCTTTTTTTCCGTAGGCAGATTTTCTGGAGATAACCTGCGCCATAAAAGATATGATGCAATGATTGAATCGAAGGATGTGCTGGAAGAGGAAGCGGTAAGTGATGTATTTCTTAATATTGTAAGCCGCTCAAATCTAAAAAGCGTAATCCTTAATTATATATGTTTAAAAACATTCTTCTTATTTATACCGTTTGCATTATGGTCACTTCGCACCGTTCTTATGCGGGAGGGATATATAAGCCGGATGTTTGTGATACGTTATATTCATAATTCCGACGGAGAAA AAGAAGCAGCAGGCAGGCTATGGCAGATTGAAAAT<u>TAATAATAATAA</u> -3’; underlined are the T7 promoter region, ribosomal binding site, start codon, FLAG-tag and stop codon, respectively). DNA oligo primers used were purchased from Metabion. PCR conditions applied were as suggested by the manufacturer and PCR products were purified via spin columns, and in vitro transcription reaction was set up using 1 μg PCR product per 50 μl reaction volume and T7 RNA polymerase (Thermo Scientific™). RNA was purified by LiCl precipitation and washed with ethanol.

### *Bacillus subtilis* S12 translation extract

Cell extracts for in vitro translation on *B. subtilis* ribosomes were generated as described previously^17,32^ but using the *B. subtilis* strain 168 *ΔyvyD ΔssrA ΔsmpB* and with some modifications in the procedure. Briefly, cells (*B. subtilis* strain 168 *ΔyvyD ΔssrA ΔsmpB*) were grown to OD600 0.8 in 2x YPTG medium (16 g L^−1^ peptone, 10 g L^−1^ yeast extract, 5 g L^−1^ NaCl, 22 mM NaH2PO4, 40 mM Na2HPO4, 19.8 g L^−1^ glucose, sterile-filtered) at 37 °C. Cells were collected by centrifugation at 5000 rpm at room temperature for 15 min and subsequently washed 3x in room temperature Buffer A (10 mM Tris–acetate (pH 8.2, 4 °C), 14 mM magnesium acetate, 60 mM potassium glutamate, 1 mM dithiothreitol, and 6 mM 2-mercaptoethanol, sterile-filtered). After the third wash, cells were resuspended in a minimal volume (0.7 mL g^−1^) of room temperature Buffer B (Buffer A without 2-mercaptoethanol). Cells were snap-frozen in liquid nitrogen and stored at −80 °C. Cells were subsequently thawed on ice and then lysed using FastPrep−24TM MP (4 × 30 sec, shaking 4.5 m s^−1^ intercalated by 1 min rest on ice), the lysate was collected by centrifugation (1000 × g, 4 °C, 1 min) and further cleared by centrifugation at 12,000 × g, 4 °C, 10 min. The lysate was used immediately, or aliquoted, snap-frozen, and stored at −80 °C.

### Generation of the complexes

To generate the LudP-SRC, the LudP mRNA template (600 ng μL^−1^) was translated by incubation in a *B. subtilis* in vitro translation system. Briefly, a total reaction volume of 400 μL was prepared by mixing 104 μL reconstitution buffer, 8 μL of methionine, 80 μL amino acid mix, 68 μL reaction mix (from RTS 100 HY Kit from Biotechrabbit GmbH) with 80 μL *B. subtilis* S12 translation extract, 4 μL LudP mRNA, 25 μL 200 mM magnesium acetate, 1 µL Murine RNase Inhibitor (NEB), and then incubated for 20 min at 30 °C shaking in a thermomixer (500 rpm).

### Purification of the complexes for structural analysis

The complex previously generated was purified by incubating the reaction with 50 μL anti-FLAG® M2 affinity gel (Merck), previously equilibrated with Hico buffer (50 mM HEPES-KOH (pH=7.4, 4 °C), 100 mM potassium acetate, 15 mM magnesium acetate, 1 mM dithiothreitol, 0.01 % (w/v) n-dodecyl-beta-maltoside, sterile-filtered) inside Mobicol column provided with 35 μm filter (MoBiTec) at 4 °C for 3 hours while rolling. After removal of the flowthrough, the beads were washed with a total of 2 mL Hico buffer and then the bound complex was eventually eluted by incubation with 15 μL Hico buffer containing 0.6 mg mL^-1^ 3xFLAG peptide for 45 min at 4 °C while rolling, followed by centrifugation (2000 g, 4 °C, 2 min). Aliquots from each fraction were checked by western blotting.

### Cryo-EM sample preparation

3.5 µl of sample (15 OD_260_/ml) were applied to grids (Quantifoil, Cu, 300 mesh, R3/3 with 3 nm carbon) which had been freshly glow discharged using a GloQube Plus (Quorum Technologies) in negative charge at 25 mA for 30 sec to make grids hydrophilic. Sample vitrification was performed using mixture of ethane/propane in 1:2 ratio in a Vitrobot Mark IV (ThermoScientific), the chamber was set to 4°C and 100% rel. humidity, blotting was done for 3 sec with zero blot force with Whatman 597 blotting paper. The grids were subsequently clipped in Autogrids cartridges and stored in liquid nitrogen until high resolution data collection.

### Cryo-EM Data collection

Data collection was performed on 300kV Titan Krios (Thermo Fisher/FEI) with Fringe-Free Imaging (FFI) setup and equipped with Gatan K3 direct electron detector using EPU (version 3.2.0.4775REL). Magnification of 105 000x was used, with data collected using super resolution counted mode at 0.415 pixel size, binned twice on the fly through EPU yielding 0.83 pixel size. Total 40 e^-^/ Å^2^ fluence was fractionated into 35 frames resulting in 1.14 e^-^/A^2^ dose per frame and total exposure of 1.91 seconds in Nanoprobe mode (15 e^-^/px/s over an empty area on the camera level). Defocus range of -0.3 to -0.9 was used with step size of 0.1 between holes. C2 aperture of 70 µm was inserted with beam spot size of 7. BioQuantum energy filter set to 20 eV cut-off was used to remove inelastically scattered electrons. Final objective astigmatism correction <1 nm and auto coma free alignment <40 nm was achieved using AutoCTF function of Sherpa (version 2.11.1). A total of 12,642 micrographs were collected for LudP-SRC (12 exposures per hole) and saved as tiff gain corrected files.

### Single-particle reconstruction of SRC complexes

RELION (version 5.0.0)^33,34^ was used for processing, unless otherwise specified. For motion correction, RELION’s implementation of MotionCor2 with 7 × 5 patches, and, for initial contrast transfer function (CTF) estimation, CTFFIND version 4.1.14^35^, were employed.

From 12,642 micrographs, a total of 1,555 micrographs with lacking CTF fit to higher spatial frequencies were excluded and 748,174 particles were picked from the remaining 11,087 micrographs using crYOLO with a general model^36^. In total, 254,695 ribosome-like particles were selected after two-dimensional (2D) classification and extracted at 4× decimated pixel size (3.324 Å per pixel) (**Supplementary Fig. 5**). An initial three-dimensional (3D) consensus refinement was done using a molmap based on *E. coli* 70S ribosome (PDB ID 7K00^37^) without mRNA and tRNAs (**Supplementary Fig. 5b**), followed by a classification with 5 classes. One class contained 70S ribosomes with A- and P-tRNA (59,787 particles), one class contained 70S ribosomes at rather low resolution (58,149 particles), one class contained 70S ribosomes with P-tRNA and substoichiometric A-tRNA (10,795 particles), on class containing 70S ribosomes with P-tRNA and a vacant A-site (125,814 particles) and one class with non-aligning particles (150 particles) (**Supplementary Fig. 5c**). All classes containing ribosome-like particles were pooled and subjected to partial signal subtraction with a mask around A- and P-sites to perform focused classification with 5 classes. This initial focus sort led to one class containing 70S ribosomes with A/P- and P/E-tRNAs (11,713 particles), one class containing 70S ribosomes with A-and P-tRNA (79,840 particles), one class containing 70S ribosomes with P-tRNA (162,522 particles) and two classes containing non-aligning particles (234 particles and 236 particles) (**Supplementary Fig. 5d**). Classes containing 70S ribosome-like particles were subsorted with a mask around the A-site to exclude non-aligning and low-resolution particles (**Supplementary Fig. 5e-g**). 155,579 particles representing a 70S ribosome with P-tRNA (**Supplementary Fig. 5e**), 11,650 particles representing a 70S ribosome with A/P- and P/E-tRNA (**Supplementary Fig. 5f**) and 79,551 particles representing a 70S ribosome with A- and P-tRNA (**Supplementary Fig. 5g**) were reverted to their original images, extracted in full resolution (0.831 Å per pixel) and 3D refined, respectively. However, only the reconstruction that contained A- and P-tRNA showed a defined nascent chain within the NPET. Therefore, only this class was CTF refined (4th order aberrations, beam tilt, anisotropic magnification and per-particle defocus value estimation), then subjected to Bayesian polishing^38^ and another round of CTF refinement. For the LudP-SRC with A-tRNA and P-tRNA a final resolution (gold-standard FSC0.143) of masked reconstructions of 2.6 Å was achieved (**Supplementary Fig. 5h; Supplementary Fig. 6**).

To estimate local resolution values, Relion’s implementation was used on one half-map of the final reconstructions (B-factor -50.0884) (**Supplementary Fig. 6e–f, h, j**).

### Molecular modeling of SRC complexes

The molecular models of the 30S and 50S ribosomal subunits were based on the B. subtilis 70S ribosome (PDB ID 9FY3^8^ for LudP-SRC. The tRNAs and nascent chains were modeled de novo. Restraint files for modified residues were created using aceDRG^39^. Starting models were rigid body fitted using ChimeraX^40^ and modelled using Coot 0.9.8.96^41^ from the CCP4 software suite version 8.0^42^. The sequence for the tRNAs was adjusted based on the appropriate anticodons corresponding to the mRNA. Final refinements were done in REFMAC 5^43^ using Servalcat^44^. The molecular models were validated using Phenix comprehensive cryo-EM validation in Phenix 1.20–4487^45^.

### Molecular Dynamics simulations

To obtain dynamics of ribosome-bound LudP at physiological temperature and to observe the effects of the D133A mutation, we carried out all-atom explicit-solvent MD simulation of the region around the LudP peptide. To that aim, all residues and ions within 35 Å of the LudP peptide were extracted from the cryo-EM model. The residues covalently bound to residues outside of the region were treated as described earlier^46^. A-tRNA atoms were removed and for the D133A variant all side chain atoms apart from Cβ were removed from Asp133. Amino acid protonation states were determined using PROPKA3^47^. The two structures were placed at the center of a dodecahedron box with a minimum distance of 15 Å between the atoms and the box boundary. The systems were solvated with the program solvate^48^ and the charge was neutralized by adding K^+^ ions using GENION^48^. Next, 7 mM MgCl_2_ and 150 mM KCl were added using GENION. All simulations were performed with GROMACS 2025^48^ using the amber14sb force field for protein^49^ and OL3 for RNA^50^. The TIP3P water model^51^, microMg parameters for Mg^2+^ ^52^, and the K^+^ and Cl^−^ parameters from ref. ^53^ were used. Short-range electrostatic and Lennard-Jones interactions were calculated for distances below 10 Å; for larger distances, long-range electrostatics were computed by particle-mesh Ewald summation^54^ with a grid spacing of 12 Å. Bond lengths were constrained using LINCS^55^ and virtual site constraints were used for hydrogens^56^, allowing an integration time step of 4 fs. Velocity rescaling^57^ was used for temperature coupling to a heat bath at T = 300 K with a coupling time constant of 5 ps, independently for solute and solvent. The pressure was controlled using stochastic cell rescaling^58^ with a reference pressure of 1 bar and a time constant of 5 ps. First, each system was energy minimized by steepest descent, applying harmonic position restraints to solute heavy atoms (k=1000 kJ mol−1 nm−2) and then the solvent was equilibrated for 50 ns keeping the position restraints. Next, position restraints were linearly decreased to zero for all residues within 25 Å of the LudP peptide during 20 ns. Subsequently, 2-µs simulations were carried out and used for the following analysis. The equilibration and production runs were repeated for 10 independent replicas per system and solute atom coordinates were recorded every 5 ps.

Prior to analysis, all trajectories were aligned to the cryo-EM model by rigid-body fitting of the rRNA P atoms. To monitor the dynamics of nucleotides U2506 and U2585, we carried out a principal component analysis (PCA)^59^ of the heavy atoms of each nucleotide separately after concatenating all trajectories with WT and D133A LudP. For each nucleotide, histograms of the projections of the trajectories onto the first two eigenvectors, i.e. conformational modes, were calculated along with the projections of the cryo-EM model and the pre-attack conformation^21^. To monitor the dynamics of the LudP peptide, we carried out a PCA of the backbone atoms of the 5 C-terminal LudP amino acids. The overlap of LudP with the pre-attack conformations of U2506 and U2585 was calculated by using a Monte-Carlo volume estimation algorithm as described previously^26^. Overlap volumes were calculated for the cryo-EM model and for trajectory frames at intervals of 500 ps.

### Figures

UCSF ChimeraX 1.10 was used to isolate density and visualize density images and structural superpositions. Models were aligned using Matchmaker function in ChimeraX 1.10. Figures were assembled with Inkscape 1.3.

## Data availability

The raw sequencing reads generated by deep mutational scanning have been deposited in the DDBJ Sequence Read Archive (DRA) under BioProject accession number PRJDB45792. Cryo-EM maps have been deposited in the Electron Microscopy Data Bank (EMDB) with accession codes EMD-58773 (LudP-SRC with A-and P-tRNA). Molecular model has been deposited in the Protein Data Bank with accession code 32BG (LudP-SRC with A- and P-tRNA). Publicly available data used included PDB ID 9MTP, 8QOA, 5NWY, 7O19, 5A81, 8CVK and 9FY1. Source data are provided with this paper.

## Code availability

Code for deep mutational scanning analysis are available on GitHub at https://github.com/KigFjwr/arrest-peptide-DMS/releases/tag/v0.3.0. MD simulation input files, final coordinates and analyses are available on zenodo at https://doi.org/10.5281/zenodo.23033215.

## Acknowledgements

We thank Machiko Murata and Naoko Muraki for their technical support. Cryo-EM data collection was performed at the Multi-User CryoEM Facility at the Centre for Structural Systems Biology, Hamburg, supported by the Universität Hamburg and DFG grant numbers (INST 152/772-1|152/774-1|152/775-1|152/776-1|152/777-1 FUGG), the Federal Ministry of Education and Research, and the DLR Projektträger (project SEEK 01KX2220). The funders had no role in study design, data collection and analysis, decision to publish, or preparation of the manuscript. We also acknowledge financial support from the Open Access Publication Fund of the Universität Hamburg.

## Author contributions

K.F. performed bioinformatics analysis, including screening to identify LudP. Y.H., K.F. and S.C. performed biochemical and genetics analyses. A.H. and F.G. prepared the LudP-SRC. H.A.S. prepared cryo-EM grids, screened and collected the cryo-EM data. F.G. performed the cryo-EM analysis, as well as generated and refined the molecular models. MD simulations were performed by L.V.B.. S.C., K.F., D.N.W., F.G. wrote the manuscript with input from all authors. S.C. and D.N.W conceived and supervised the project.

