## Supplemental Information for "The LudP arrest peptide stalls translation by remodeling the ribosomal A-site"

Supplementary Information
for

**The LudP arrest peptide stalls translation by remodeling the ribosomal A-site**

Felix Gersteuer<sup>1,\*</sup>, Yuki Harada<sup>2,\*</sup>, Keigo Fujiwara<sup>2,3</sup>, Haaris A. Safdari<sup>1</sup>, Alena Harreither<sup>1</sup>, Lars V. Bock<sup>4</sup>, Daniel N. Wilson<sup>1,#</sup>, Shinobu Chiba<sup>2,#</sup>

<sup>1</sup> Institute for Biochemistry and Molecular Biology, Martin-Luther-King-Platz 6, University of Hamburg, 20146 Hamburg, Germany.

<sup>2</sup> Faculty of Life Sciences and Institute for Protein Dynamics, Kyoto Sangyo University, Kamigamo, Motoyama, Kita-ku, Kyoto 603-8555, Japan.

<sup>3</sup> Department of Gene Function and Phenomics, National Institute of Genetics, Mishima, Japan

<sup>4</sup> Department of Theoretical and Computational Biophysics, Max Planck Institute for Multidisciplinary Sciences, Göttingen, Germany

\* These authors contributed equally

### Corresponding authors:

Prof Shinobu Chiba

Prof Daniel N. Wilson

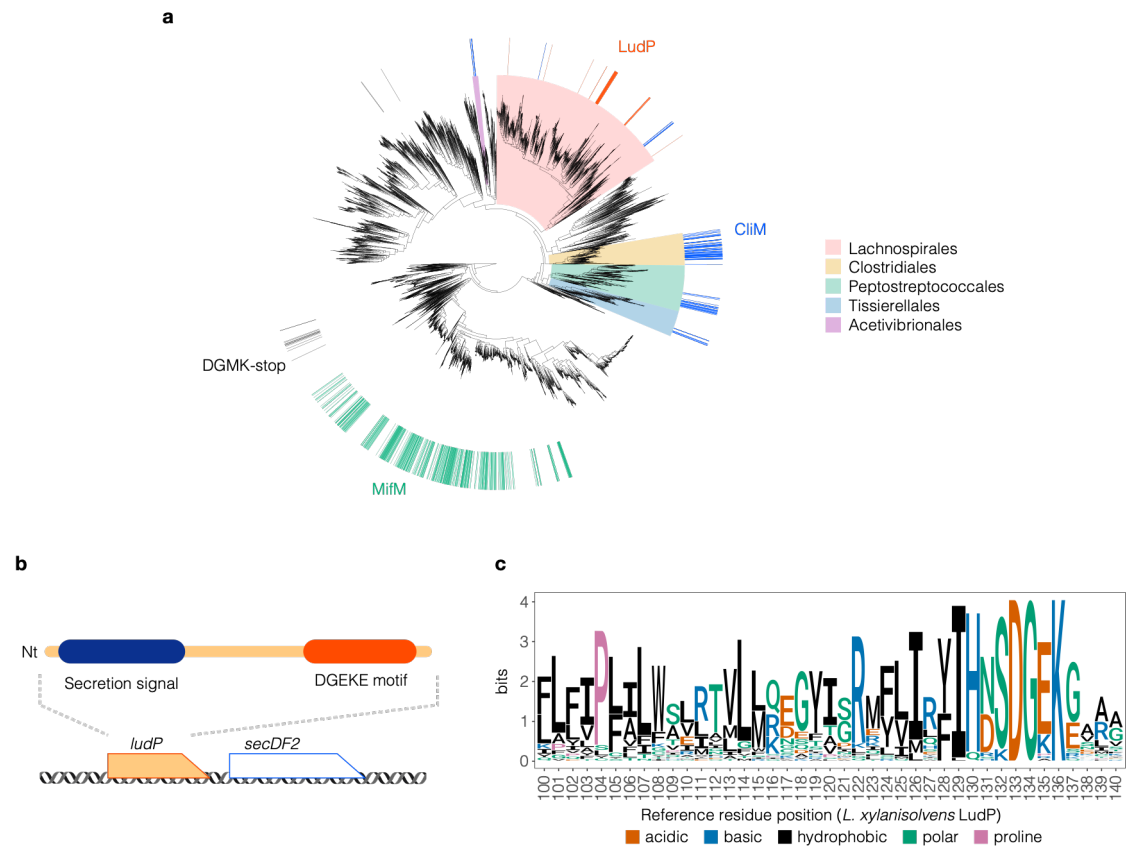

**Supplementary Fig. 1: Sequence conservation and phylogenetic distribution of LudP homologs.**

**a.** Distribution of LudP and other arrest peptides on phylum Bacillota. Colored outer marks indicate genomes encoding LudP (vermilion), MifM (green), CliM (blue), or uDF-DGMK-stop (gray). Background shading highlights five orders within the class Clostridia. **b.** Schematic of *Lacrimispora xyloisolvans* LudP and its genomic context. **c.** Sequence logo of 49 LudP homologs spanning the alignment region corresponding to residues 100-140 of *L. xyloisolvans* LudP.

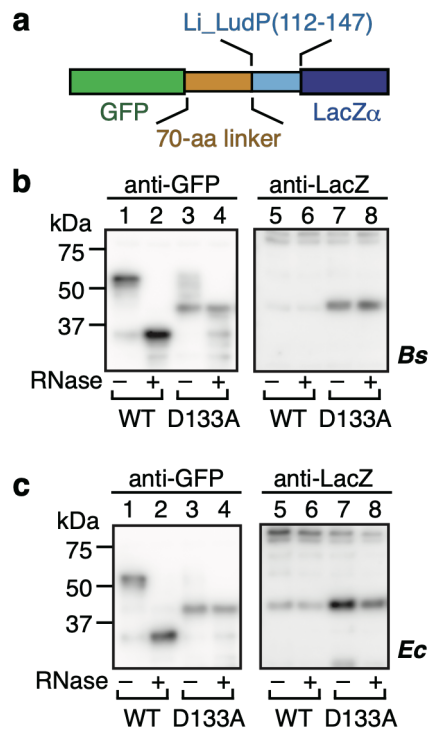

**Supplementary Fig. 2: *In vitro* translation assay of *L. indoles* LudP.** **a.** Schematic representation of the *gfp-Li\_ludP(112-147)-lacZα* reporter used for *in vitro* translation assay. **b, c.** *In vitro* translation assay. *gfp-Li\_ludP(112-147)-lacZα* was translated in Bs PURE (**b**) or Ec PURE (**c**) at 37°C for 20 min. Translation products were analyzed by immunoblotting with anti-GFP (left) and anti-LacZ (right) antibodies. A portion of each sample was treated with RNase A to digest the tRNA moiety of peptidyl-tRNA.

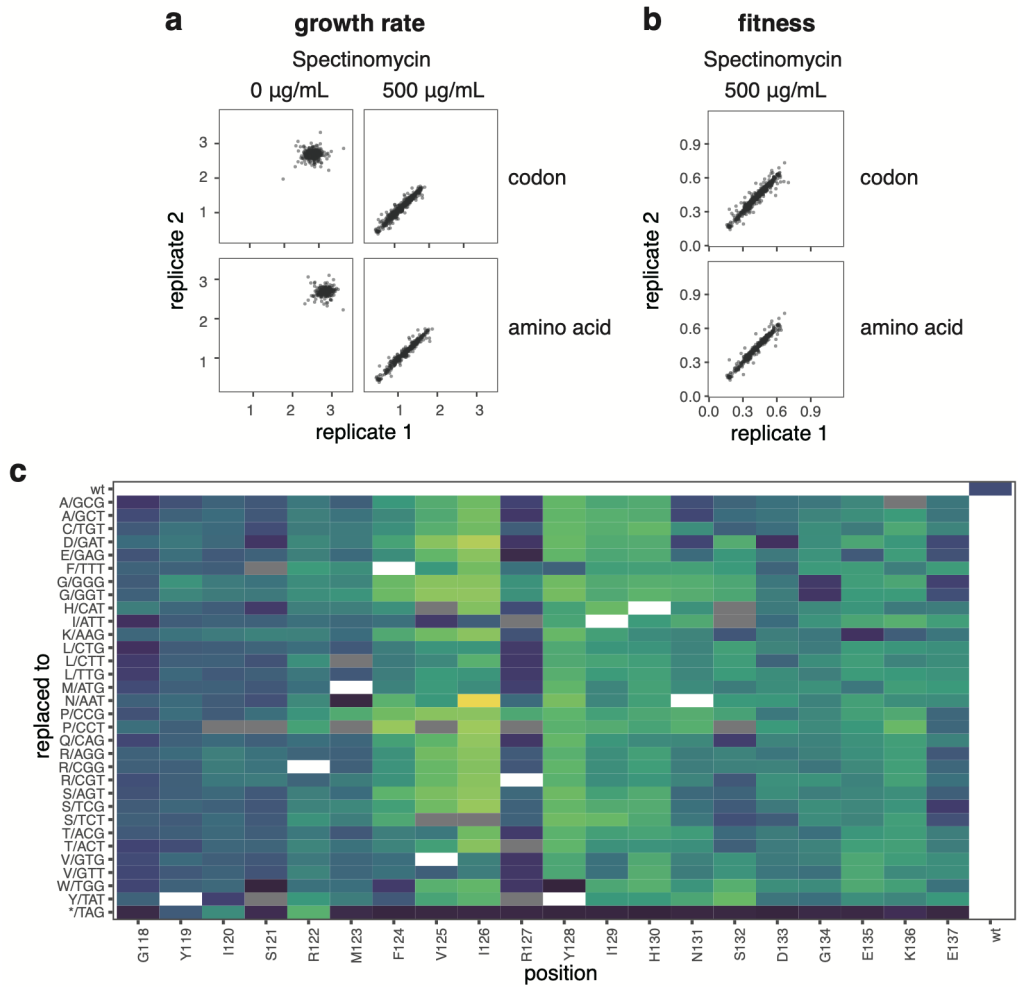

**Supplementary Fig. 3: Reproducibility of deep mutational scanning of LudP. a, b.** Scatter plots of growth rates (a) and fitness values (b) for each variant are shown at the codon level (upper) and at the amino acid level (lower). Amino acid-level values were obtained by averaging the values of synonymous codon variants. In each plot, biological replicate 1 is shown on the x-axis and biological replicate 2 on the y-axis, showing good reproducibility between replicates. **c.** Heatmap showing the relative fitness of each variant at the codon level, calculated as the mean of two biological replicates. Synonymous codon variants generally exhibited similar fitness values. Residue numbers and wild-type amino acids are indicated along the bottom, and substituted amino acids and their corresponding codons are shown along the left.

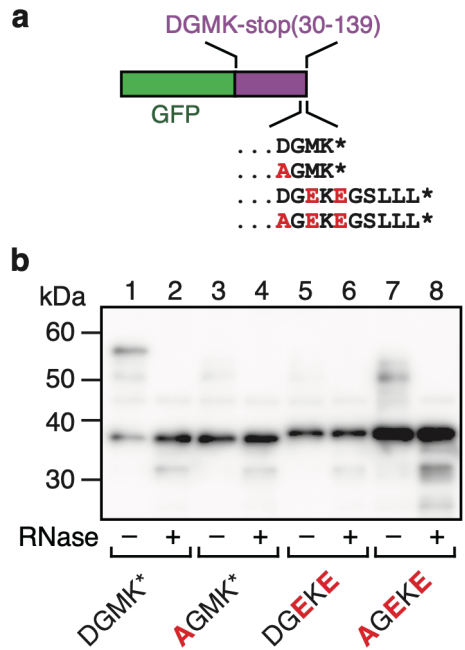

**Supplementary Fig. 4: *In vitro* translation assay of uDF\_DGMK-stop variants containing LudP-derived C-terminal sequences.** **a.** Schematic representation of the *gfp-uDF\_DGMK-stop(112-147)-lacZα* reporter and the C-terminal amino acid sequences of the variants used for the *in vitro* translation assay. **b.** *In vitro* translation assay. *gfp-uDF\_DGMK-stop(112-147)-lacZα* was translated in Bs PURE at 37°C for 20 min. Translation products were analyzed by immunoblotting with an anti-GFP antibody. A portion of each sample was treated with RNase A to digest the tRNA moiety of peptidyl-tRNA. Only the wild-type reporter (DGMK\*) produced an RNase-sensitive peptidyl-tRNA product that migrated between the 50- and 60-kDa markers.

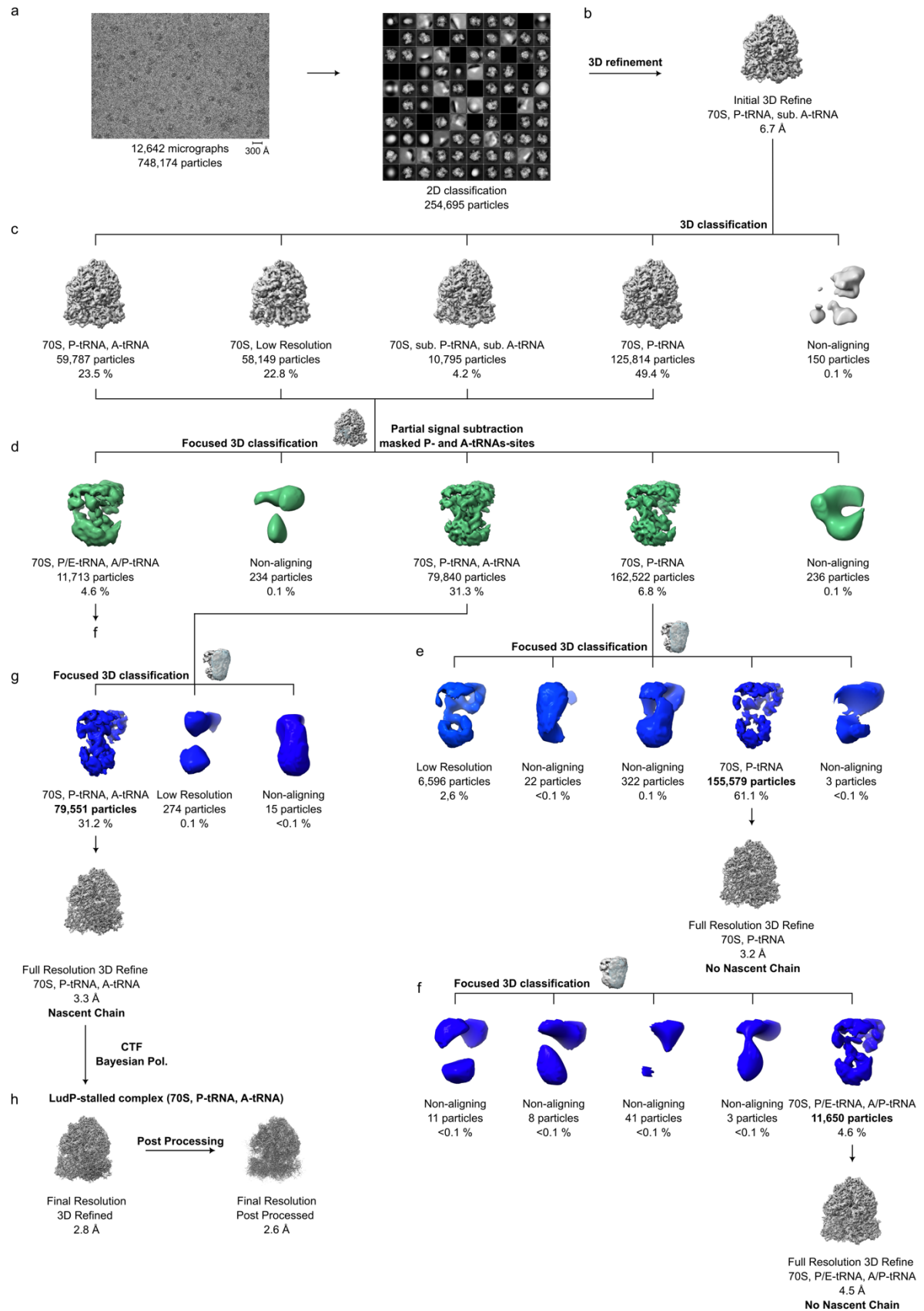

**Supplementary Fig. 5: *In silico* sorting scheme for LudP.** (a) From 12,642 micrographs, 741,174 particles were picked and subjected to 2D classification resulting in 254,695 ribosome-like particles. Particles were (b) initially 3D-refined, then (c) subsorted into 5 classes. (d) Classes containing 70S ribosome-like particle were joined and subsorted using a mask around the A- and P-tRNA binding sites. (e-g) The resulting classes with A/P- and P/E-tRNA, P- and A-tRNA as well as P-tRNA and vacant A-site were further subsorted with a mask around the A-site until homogeneity was reached. The minor class with A/P- and P/E-tRNA (4,6 %) (f) as well as the major class with P-tRNA and vacant A-site (61.1%) (e) were refined in full resolution to 4.5 Å and 3.2 Å (at FSC 0.143), respectively. However, no stable nascent peptide was detectable within the NPET. The median class with A- and P-tRNA (31.2 %) revealed a stable nascent peptide in the NPET and resulted in a final resolution (at FSC 0.143) of 2.6 Å.

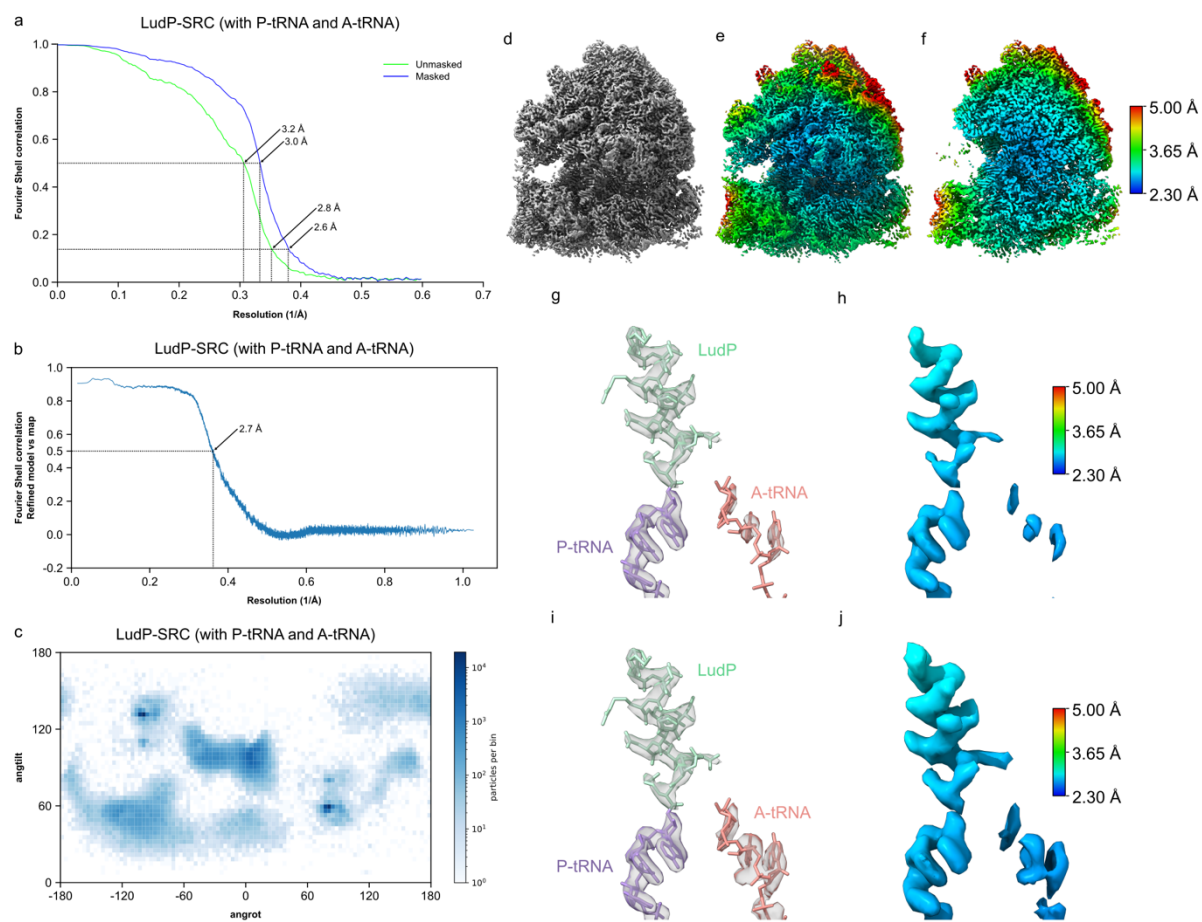

**Supplementary Fig. 6: Fourier Shell Correlation curves, angular distribution and local resolution for LudP-SRC.** (a) Fourier shell correlation (FSC) curve of the LudP-SRC containing A- and P-tRNA with unmasked (green) and masked (blue) FSC curves plotted against the resolution (1/Å). (b) Refined model vs map FSC curve of the LudP-SRC from (a) plotted against the resolution (1/Å). (c) Angular distribution of particles used for 3D reconstruction from Relion for the LudP-SRC from (a). Particles are binned and logarithmically represented from white to blue. (d-j) Cryo-EM density for the 3D-refined map of the LudP-SRC with A- and P-tRNA in grey (d), coloured according to local resolution (e-f,h,j) or as transparent grey surface (g,i) with molecular model of LudP-SRC. In (d-e) an overview of the cryo-EM map of the LudP-SRC is shown, whereas in (f) a transverse section reveals the core of the 50S subunit, including the ribosomal exit tunnel. In (i-j) the same representation is shown as in (g-h) but at a lower threshold.

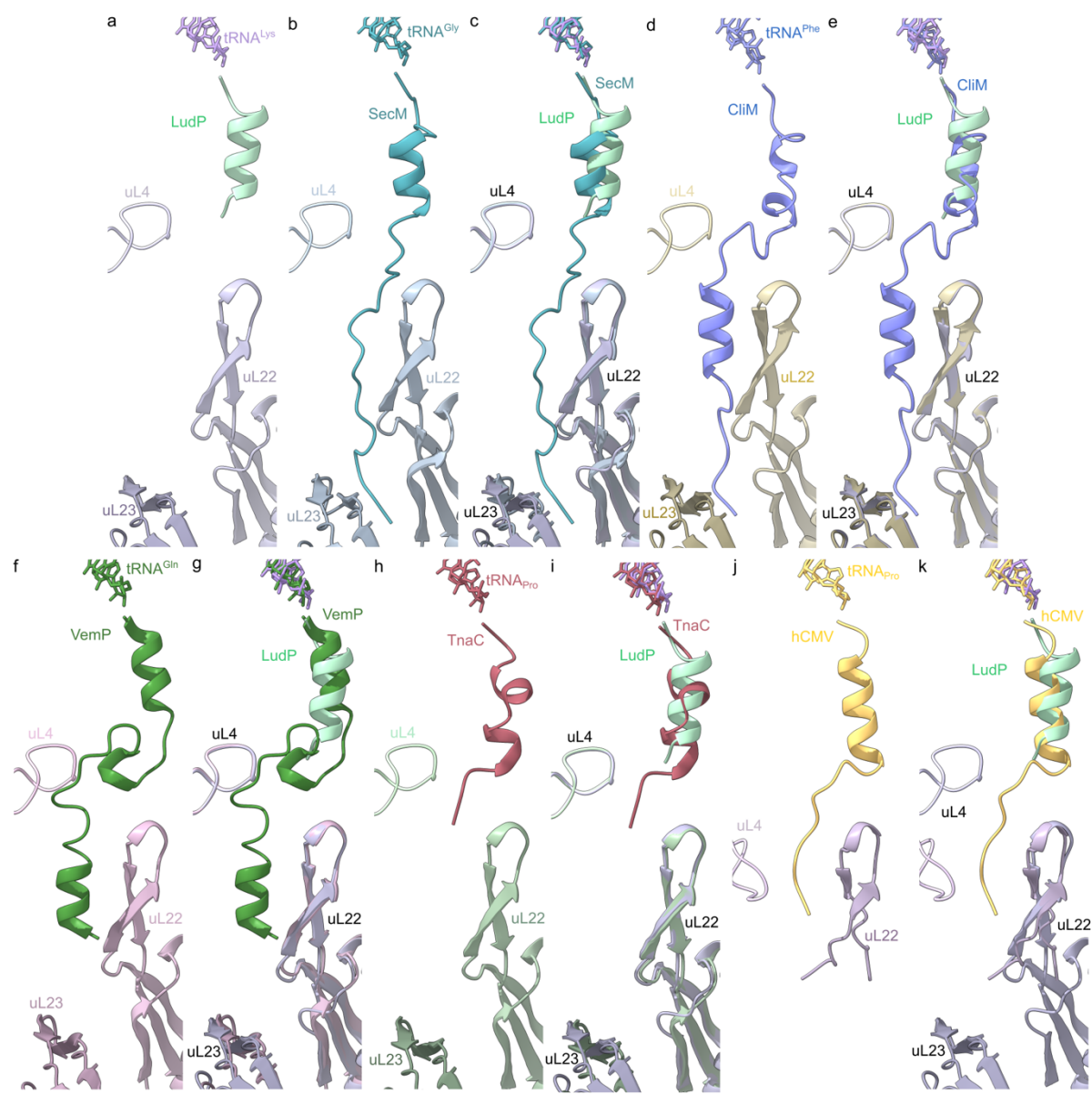

**Supplementary Fig. 7: Comparison of LudP with SecM, CliM, VemP, TnaC and hCMV arrest peptides.** (a) LudP (mint) attached to the P-tRNA (lavender) in relation to uL4 (light grey), uL22 (grey) and uL23 (dark grey). (b) SecM (PDB ID 8QOA)<sup>1</sup> attached to the P-tRNA (teal) in relation to uL4 (light slate blue), uL22 (slate blue) and uL23 (dark slate blue). (c) Overlay (aligned on the basis of 23S rRNA) of (a) LudP and (b) SecM. (d) CliM (PDB ID 9FY1)<sup>2</sup> attached to the P-tRNA (blue) in relation to uL4 (light gold), uL22 (gold) and uL23 (dark gold). (e) Overlay (aligned on the basis of 23S rRNA) of (a) LudP and (d) CliM. (f) VemP (PDB ID 5NWY)<sup>3</sup> attached to the P-tRNA (green) in relation to uL4 (light rose), uL22 (rose) and uL23 (dark rose). (g) Overlay (aligned on the basis of 23S rRNA) of (a) LudP and (f) VemP. (h) TnaC (PDB ID 7O19)<sup>4</sup> attached to the P-tRNA (red) in relation to uL4 (light mint), uL22 (mint) and uL23 (dark mint). (i) Overlay (aligned on the basis of 23S rRNA) of (a) LudP and (h) TnaC. (j) hCMV (PDB ID 5A8I)<sup>5</sup> attached to the P-tRNA (gold) in relation to uL4 (pink) and uL22 (purple). (k) Overlay (aligned on the basis of 23S rRNA) of (a) LudP and (j) hCMV.

**Supplementary Table 1: Bacterial strains used in this study**

| Strain | Genotype | Host | Plasmid | Organism |
| --- | --- | --- | --- | --- |
| HRE1 | *gfp-Lx_ludP(16-147)-myc-lacZ* | JM109 | pHR9 | *E. coli* |
| HRE3 | *gfp-Lx_ludP(16-147)-D133A-myc-lacZ* | JM109 | pKIG1527 | *E. coli* |
| KFE701 | *gfp-apdP(34-103)-Li_ludP(112-146)-myc-lacZ* | JM109 | pKIG1336 | *E. coli* |
| KFE703 | *gfp-apdP(34-103)-Li_ludP(112-146)-D133A-myc-lacZ* | JM109 | pKIG1523 | *E. coli* |
| HRB1 | *amyE::PmifM gfp-Lx_ludP(16-147)-myc-lacZøcat* | PY79 | pHR9 | *B. subtilis* |
| HRB3 | *amyE::PmifM gfp-Lx_ludP(16-147)-D133A-myc-lacZøcat* | PY79 | pKIG1527 | *B. subtilis* |
| HRB89 | *amyE::PmifM SD(kf)-Lx_ludP-malF-myc-lacZøcat* | PY79 | pHR130 | *B. subtilis* |
| HRB90 | *amyE::PmifM SD(kf)-Lx_ludP-D133A-malF-myc-lacZøcat* | PY79 | pHR132 | *B. subtilis* |
| HRB118 | *amyE::PmifM SD(kf)-Lx_ludP(Δ6-25)-malF-myc-lacZøcat* | PY79 | pHR172 | *B. subtilis* |
| HRB119 | *amyE::PmifM SD(kf)-Lx_ludP(Δ6-25)-D133A-malF-myc-lacZøcat* | PY79 | pHR174 | *B. subtilis* |
| HRB123 | *amyE::PmifM gfp-Lx_ludP(16-147)-FS15-myc-lacZøcat* | PY79 | pCH2855 | *B. subtilis* |
| HRB122 | *amyE::PmifM gfp-Lx_ludP(16-147)-FS20-myc-lacZøcat* | PY79 | pCH2856 | *B. subtilis* |
| HRB121 | *amyE::PmifM gfp-Lx_ludP(16-147)-FS25-myc-lacZøcat* | PY79 | pCH2847 | *B. subtilis* |
| HRB120 | *amyE::PmifM gfp-Lx_ludP(16-147)-FS30-myc-lacZøcat* | PY79 | pCH2846 | *B. subtilis* |
| HRB126 | *amyE::PmifM gfp-Lx_ludP(16-147)-myc-lacZøcat, rplWøkan* | SCB2613 | pHR9 | *B. subtilis* |
| HRB127 | *amyE::PmifM gfp-Lx_ludP(16-147)-myc-lacZøcat, rplD(Δ66-70)økan* | SCB2942 | pHR9 | *B. subtilis* |
| HRB130 | *amyE::PmifM gfp-Lx_ludP(16-147)-D133A-myc-lacZøcat, rplWøkan* | SCB2613 | pHR54 | *B. subtilis* |
| HRB131 | *amyE::PmifM gfp-Lx_ludP(16-147)-D133A-myc-lacZøcat, rplD(Δ66-70)økan* | SCB2942 | pHR54 | *B. subtilis* |
| HRB69 | *amyE::PmifM gfp-Lx_ludP(16-147)-myc-lacZøcat, rpsSøkan* | SCB2656 | pHR9 | *B. subtilis* |
| HRB70 | *amyE::PmifM gfp-Lx_ludP(16-147)-myc-lacZøcat, rpsSøkanørplV(Δ86-90)* | SCB2917 | pHR9 | *B. subtilis* |
| HRB74 | *amyE::PmifM gfp-Lx_ludP(16-147)-D133A-myc-lacZøcat, rpsSøkan* | SCB2656 | pHR54 | *B. subtilis* |
| HRB75 | *amyE::PmifM gfp-Lx_ludP(16-147)-D133A-myc-lacZøcat, rpsSøkanørplV(Δ86-90)* | SCB2917 | pHR54 | *B. subtilis* |
| HRB71 | *amyE::PmifM gfp-Lx_ludP(16-147)-myc-lacZøcat, rplWøkan* | SCB2613 | pHR9 | *B. subtilis* |
| HRB72 | *amyE::PmifM gfp-Lx_ludP(16-147)-myc-lacZøcat, rplW(Δ65-69)økan* | SCB2634 | pHR9 | *B. subtilis* |
| HRB76 | *amyE::PmifM gfp-Lx_ludP(16-147)-D133A-myc-lacZøcat, rplWøkan* | SCB2613 | pHR54 | *B. subtilis* |
| HRB77 | *amyE::PmifM gfp-Lx_ludP(16-147)-D133A-myc-lacZøcat, rplW(Δ65-69)økan* | SCB2634 | pHR54 | *B. subtilis* |
| HRB105 | *amyE::PmifM gfp-Lx_ludP(16-147)-S121A-myc-lacZøcat* | PY79 | pHR148 | *B. subtilis* |
| HRB106 | *amyE::PmifM gfp-Lx_ludP(16-147)-R122A-myc-lacZøcat* | PY79 | pHR150 | *B. subtilis* |
| HRB107 | *amyE::PmifM gfp-Lx_ludP(16-147)-M123A-myc-lacZøcat* | PY79 | pHR151 | *B. subtilis* |
| HRB108 | *amyE::PmifM gfp-Lx_ludP(16-147)-F124A-myc-lacZøcat* | PY79 | pHR152 | *B. subtilis* |
| HRB109 | *amyE::PmifM gfp-Lx_ludP(16-147)-V125A-myc-lacZøcat* | PY79 | pHR154 | *B. subtilis* |
| HRB110 | *amyE::PmifM gfp-Lx_ludP(16-147)-I126A-myc-lacZøcat* | PY79 | pHR156 | *B. subtilis* |
| HRB111 | *amyE::PmifM gfp-Lx_ludP(16-147)-R127A-myc-lacZøcat* | PY79 | pHR158 | *B. subtilis* |
| HRB112 | *amyE::PmifM gfp-Lx_ludP(16-147)-Y128A-myc-lacZøcat* | PY79 | pHR160 | *B. subtilis* |
| HRB113 | *amyE::PmifM gfp-Lx_ludP(16-147)-I129A-myc-lacZøcat* | PY79 | pHR162 | *B. subtilis* |
| HRB114 | *amyE::PmifM gfp-Lx_ludP(16-147)-H130A-myc-lacZøcat* | PY79 | pHR164 | *B. subtilis* |
| HRB2 | *amyE::PmifM gfp-Lx_ludP(16-147)-N131A-myc-lacZøcat* | PY79 | pKIG1526 | *B. subtilis* |
| HRB115 | *amyE::PmifM gfp-Lx_ludP(16-147)-S132A-myc-lacZøcat* | PY79 | pHR166 | *B. subtilis* |
| HRB3 | *amyE::PmifM gfp-Lx_ludP(16-147)-D133A-myc-lacZøcat* | PY79 | pKIG1527 | *B. subtilis* |
| HRB4 | *amyE::PmifM gfp-Lx_ludP(16-147)-G134A-myc-lacZøcat* | PY79 | pKIG1528 | *B. subtilis* |
| HRB5 | *amyE::PmifM gfp-Lx_ludP(16-147)-E135A-myc-lacZøcat* | PY79 | pKIG1529 | *B. subtilis* |
| HRB6 | *amyE::PmifM gfp-Lx_ludP(16-147)-K136A-myc-lacZøcat* | PY79 | pKIG1530 | *B. subtilis* |
| HRB7 | *amyE::PmifM gfp-Lx_ludP(16-147)-E137A-myc-lacZøcat* | PY79 | pKIG1531 | *B. subtilis* |
| HRB152 | *amyE::PmifM gfp-Lx_ludP(16-147)-E137Y-myc-lacZøcat* | PY79 | pCH2903 | *B. subtilis* |
| HRB153 | *amyE::PmifM gfp-Lx_ludP(16-147)-E137F-myc-lacZøcat* | PY79 | pCH2904 | *B. subtilis* |
| HRB154 | *amyE::PmifM gfp-Lx_ludP(16-147)-E137R-myc-lacZøcat* | PY79 | pCH2905 | *B. subtilis* |
| HRB155 | *amyE::PmifM gfp-Lx_ludP(16-147)-E137D-myc-lacZøcat* | PY79 | pCH2907 | *B. subtilis* |
| HRB146 | *amyE::PmifM gfp-Lx_ludP(16-147)-mifM91-yidC2(6)-lacZøcat* | PY79 | pCH2871 | *B. subtilis* |
| HRB147 | *amyE::PmifM gfp-Lx_ludP(16-147)-D133A-mifM91-yidC2(6)-lacZøcat* | PY79 | pCH2872 | *B. subtilis* |
| HRB148 | *amyE::PmifM gfp-Lx_ludP(16-136)\*-mifM91-yidC2(6)-lacZøcat* | PY79 | pCH2873 | *B. subtilis* |
| HRB149 | *amyE::PmifM gfp-Lx_ludP(16-136)\*-D133A-mifM91-yidC2(6)-lacZøcat* | PY79 | pCH2874 | *B. subtilis* |
| HRB150 | *amyE::PmifM gfp-Lx_ludP(16-147)(DGMK\*)-mifM91-yidC2(6)-lacZøcat* | PY79 | pCH2895 | *B. subtilis* |
| HRB151 | *amyE::PmifM gfp-Lx_ludP(16-147)(AGMK\*)-mifM91-yidC2(6)-lacZøcat* | PY79 | pCH2896 | *B. subtilis* |

Supplementary Table 2: Plasmids used in this study

| Plasmid | Genotype | Primer fw 1 | Primer rv 1 | Template 1 | Primer fw 2 | Primer rv 2 | Template |
| --- | --- | --- | --- | --- | --- | --- | --- |
| pHR9 | amyE::PmiFM gfp-Lx_ludP(16-147)-myc-lacZocat | myc24fw | GFP238 24rv | pKIG1336 | gfp-uDF-DGEKG-Lxyl-8953-fw | uDF-DGEKG-Lxyl-9302-myc-rv | uDGEKG_Lxyl |
| pHR54 | amyE::PmiFM gfp-Lx_ludP(16-147)-D133A-myc-lacZocat | ampR 121-128(Tm62) | DGEK_Lxyl D133A rv | pHR9 | DGEK_Lxyl D133A fw | ampR 121-128(Tm62).antisense | pHR9 |
| pHR62 | amyE::PmiFM SD(miFM)-Lx_ludP-malF-myc-lacZocat | Lxyl-DGEKE-miFM SD-L1-M-fw | Lxyl-DGEKE-miFM SD-L1-M-rv | pHR41 | - | - | - |
| pHR64 | amyE::PmiFM SD(miFM)-Lx_ludP-D133A-malF-myc-lacZocat | Lxyl-DGEKE-miFM SD-L1-M-fw | Lxyl-DGEKE-miFM SD-L1-M-rv | pHR42 | - | - | - |
| pHR130 | amyE::PmiFM SD(kf)-Lx_ludP-malF-myc-lacZocat | Lxyl-DGEKE-kfSD 31-fw | PmiFM-kfSD 24-rv | pHR62 | - | - | - |
| pHR132 | amyE::PmiFM SD(kf)-Lx_ludP-D133A-malF-myc-lacZocat | Lxyl-DGEKE-kfSD 31-fw | PmiFM-kfSD 24-rv | pHR64 | - | - | - |
| pHR148 | amyE::PmiFM gfp-Lx_ludP(16-147)-SI21A-myc-lacZocat | Lxyl-DGEKE-SI21A-fw | Lxyl-DGEKE-I120A-SI21A-rv | pHR9 | - | - | - |
| pHR150 | amyE::PmiFM gfp-Lx_ludP(16-147)-R122A-myc-lacZocat | Lxyl-DGEKE-R122A-fw | Lxyl-DGEKE-R122A-M123A-rv | pHR9 | - | - | - |
| pHR151 | amyE::PmiFM gfp-Lx_ludP(16-147)-M123A-myc-lacZocat | ampR 121-128(Tm62) | Lxyl-DGEKE-R122A-M123A-rv | pHR9 | Lxyl-DGEKE-M123A-fw | ampR 121-128(Tm62).antisense | pHR9 |
| pHR152 | amyE::PmiFM gfp-Lx_ludP(16-147)-F124A-myc-lacZocat | Lxyl-DGEKE-F124A-fw | Lxyl-DGEKE-F124-V125A-rv | pHR9 | - | - | - |
| pHR154 | amyE::PmiFM gfp-Lx_ludP(16-147)-V125A-myc-lacZocat | Lxyl-DGEKE-V125A-fw | Lxyl-DGEKE-F124-V125A-rv | pHR9 | - | - | - |
| pHR156 | amyE::PmiFM gfp-Lx_ludP(16-147)-I126A-myc-lacZocat | Lxyl-DGEKE-I126A-fw | Lxyl-DGEKE-I126A-R127A-rv | pHR9 | - | - | - |
| pHR158 | amyE::PmiFM gfp-Lx_ludP(16-147)-R127A-myc-lacZocat | Lxyl-DGEKE-R127A-fw | Lxyl-DGEKE-I126A-R127A-rv | pHR9 | - | - | - |
| pHR160 | amyE::PmiFM gfp-Lx_ludP(16-147)-Y128A-myc-lacZocat | Lxyl-DGEKE-Y128A-fw | Lxyl-DGEKE-Y128A-I129A-rv | pHR9 | - | - | - |
| pHR162 | amyE::PmiFM gfp-Lx_ludP(16-147)-I129A-myc-lacZocat | Lxyl-DGEKE-I129A-fw | Lxyl-DGEKE-Y128A-I129A-rv | pHR9 | - | - | - |
| pHR164 | amyE::PmiFM gfp-Lx_ludP(16-147)-H130A-myc-lacZocat | Lxyl-DGEKE-H130A-fw | Lxyl-DGEKE-H130A-rv | pHR9 | - | - | - |
| pHR166 | amyE::PmiFM gfp-Lx_ludP(16-147)-SI32A-myc-lacZocat | Lxyl-DGEKE-SI32A-fw | Lxyl-DGEKE-SI32A-rv | pHR9 | - | - | - |
| pHR172 | amyE::PmiFM SD(kf)-Lx_ludP(Δ6-25)-malF-myc-lacZocat | Lxyl-DGEKE-ss6-25aa-fw2 | Lxyl-DGEKE-ss6-25aa-rv2 | pHR130 | - | - | - |
| pHR174 | amyE::PmiFM SD(kf)-Lx_ludP(Δ6-25)-D133A-malF-myc-lacZocat | Lxyl-DGEKE-ss6-25aa-fw2 | Lxyl-DGEKE-ss6-25aa-rv2 | pHR132 | - | - | - |
| pCH2845 | amyE::PmiFM gfp-ludP(16-147)-98FS-myc-lacZocat | a63-Lxyl-98ins-fw | a64-Lxyl-K97-rv | pHR9 | - | - | - |
| pCH2846 | amyE::PmiFM gfp-Lx_ludP(16-147)-FS30-myc-lacZocat | a71-Lxyl-FS30-fw | a64-Lxyl-K97-rv | pHR9 | a63-Lxyl-98ins-fw | a72-Lxyl-FS30-rv | pHR9 |
| pCH2847 | amyE::PmiFM gfp-Lx_ludP(16-147)-FS25-myc-lacZocat | a69-Lxyl-FS25-fw | a64-Lxyl-K97-rv | pHR9 | a63-Lxyl-98ins-fw | a70-Lxyl-FS25-rv | pHR9 |
| pCH2855 | amyE::PmiFM gfp-Lx_ludP(16-147)-FS15-myc-lacZocat | a65-Lxyl-FS15-fw | a66-Lxyl-FS15-rv | pCH2845 | - | - | - |
| pCH2856 | amyE::PmiFM gfp-Lx_ludP(16-147)-FS20-myc-lacZocat | a67-Lxyl-FS20-fw | a64-Lxyl-K97-rv | pHR9 | a63-Lxyl-98ins-fw | a68-Lxyl-FS20-rv | pHR9 |
| pCH2865 | amyE::PmiFM gfp-uSecDF_DGMK*(30-139)*-miFM91-yidC2(6)-lacZocat | b27-miFM91-fw | gfp238-rv | pCH835 | gfp238-fw | b36-DGMK-miFM91-rv | pCH2400 |
| pCH2866 | amyE::PmiFM gfp-uSecDF_DGMK*(30-139)*-D136A-miFM91-yidC2(6)-lacZocat | b27-miFM91-fw | gfp238-rv | pCH835 | gfp238-fw | b37-DGMK-D136A-miFM91-rv | pCH2418 |
| pCH2871 | amyE::PmiFM gfp-Lx_ludP(16-147)-miFM91-yidC2(6)-lacZocat | b27-miFM91-fw | gfp238-rv | pCH835 | gfp238-fw | b42-Lxyl_DGEKE-miFM91-rv | pHR9 |
| pCH2872 | amyE::PmiFM gfp-Lx_ludP(16-147)-D133A-miFM91-yidC2(6)-lacZocat | b27-miFM91-fw | gfp238-rv | pCH835 | gfp238-fw | b43-Lxyl_AGEKE-miFM91-rv | pKIG1527 |
| pCH2873 | amyE::PmiFM gfp-Lx_ludP(16-136)*-miFM91-yidC2(6)-lacZocat | b27-miFM91-fw | gfp238-rv | pCH835 | gfp238-fw | b44-Lxyl_DGEK-ST-miFM91-rv | pHR9 |
| pCH2874 | amyE::PmiFM gfp-Lx_ludP(16-136)*-D133A-miFM91-yidC2(6)-lacZocat | b27-miFM91-fw | gfp238-rv | pCH835 | gfp238-fw | b45-Lxyl_AGEK-ST-miFM91-rv | pKIG1527 |
| pCH2895 | amyE::PmiFM gfp-Lx_ludP(16-147)(DGMK*)-miFM91-yidC2(6)-lacZocat | b27-miFM91-fw | gfp238-rv | pCH835 | gfp238-fw | b86-Lxyl_AGMK-ST-miFM91-rv | pHR9 |
| pCH2896 | amyE::PmiFM gfp-Lx_ludP(16-147)(AGMK*)-miFM91-yidC2(6)-lacZocat | b27-miFM91-fw | gfp238-rv | pCH835 | gfp238-fw | b45-Lxyl_AGEK-ST-miFM91-rv | pKIG1527 |
| pCH2899 | amyE::PmiFM rbsm1 gfp-uSecDF_DGMK*(30-139)(DGEKE)-miFM91-yidC2(6)-lacZocat | b27-miFM91-fw | gfp238-rv | pCH835 | gfp238-fw | b89-DGMK_DGEKE-miFM91-rv | pCH2400 |
| pCH2900 | amyE::PmiFM rbsm1 gfp-uSecDF_DGMK*(30-139)(AGEKE)-miFM91-yidC2(6)-lacZocat | b27-miFM91-fw | gfp238-rv | pCH835 | gfp238-fw | b90-DGMK_AGEKE-miFM91-rv | pCH2418 |
| pCH2903 | amyE::PmiFM gfp-Lx_ludP(16-147)-E137Y-myc-lacZocat | b93-DGEK-FY-fw | b96-DGEKE-136-rv | pHR9 | - | - | - |
| pCH2904 | amyE::PmiFM gfp-Lx_ludP(16-147)-E137F-myc-lacZocat | b93-DGEK-FY-fw | b96-DGEKE-136-rv | pHR9 | - | - | - |
| pCH2905 | amyE::PmiFM gfp-Lx_ludP(16-147)-E137R-myc-lacZocat | b94-DGEK-IR-fw | b96-DGEKE-136-rv | pHR9 | - | - | - |
| pCH2907 | amyE::PmiFM gfp-Lx_ludP(16-147)-E137D-myc-lacZocat | b95-DGEK-DG-fw | b96-DGEKE-136-rv | pHR9 | - | - | - |
| pKIG1089 | amyE::PmiFM gfp-Sm_apdP(34-103)-Bs_yweI(61-96)-myc-lacZ | - | - | - | - | - | - |
| pKIG1336 | amyE::PmiFM gfp-apdP(34-103)-Li_ludP(112-146)-myc-lacZocat | 5_uD_Lind_DGEKG fw | ampR 121-128(Tm62).antisense | pKIG1089 | ampR 121-128(Tm62) | 5_uD_Lind_DGEKG rv | pKIG1089 |
| pKIG1523 | amyE::PmiFM gfp-apdP(34-103)-Li_ludP(112-146)-D133A-myc-lacZocat | ampR 121-128(Tm62) | DGEKG_Lind H130 rv | pKIG1336 | DGEKG_Lind D133A fw | ampR 121-128(Tm62).antisense | pKIG1336 |
| pKIG1526 | amyE::PmiFM gfp-Lx_ludP(16-147)-N131A-myc-lacZocat | ampR 121-128(Tm62) | DGEK_Lxyl N131A rv | pHR9 | DGEK_Lxyl N131A fw | ampR 121-128(Tm62).antisense | pHR9 |
| pKIG1527 | amyE::PmiFM gfp-Lx_ludP(16-147)-D133A-myc-lacZocat | ampR 121-128(Tm62) | DGEK_Lxyl D133A rv | pHR9 | DGEK_Lxyl D133A fw | ampR 121-128(Tm62).antisense | pHR9 |
| pKIG1528 | amyE::PmiFM gfp-Lx_ludP(16-147)-G134A-myc-lacZocat | ampR 121-128(Tm62) | DGEK_Lxyl G134A rv | pHR9 | DGEK_Lxyl G134A fw | ampR 121-128(Tm62).antisense | pHR9 |
| pKIG1529 | amyE::PmiFM gfp-Lx_ludP(16-147)-E135A-myc-lacZocat | ampR 121-128(Tm62) | DGEK_Lxyl E135A rv | pHR9 | DGEK_Lxyl E135A fw | ampR 121-128(Tm62).antisense | pHR9 |
| pKIG1530 | amyE::PmiFM gfp-Lx_ludP(16-147)-K136A-myc-lacZocat | ampR 121-128(Tm62) | DGEK_Lxyl K136A rv | pHR9 | DGEK_Lxyl K136A fw | ampR 121-128(Tm62).antisense | pHR9 |
| pKIG1531 | amyE::PmiFM gfp-Lx_ludP(16-147)-E137A-myc-lacZocat | ampR 121-128(Tm62) | DGEK_Lxyl E137A rv | pHR9 | DGEK_Lxyl E137A fw | ampR 121-128(Tm62).antisense | pHR9 |

Supplementary Table 3: Primers used in this study

| Primer name | DNA sequence (5' - 3') |
| --- | --- |
| ampR 121-128(Tm62) | GCAGTGCTGCCATAACCATGAGTG |
| ampR 121-128(Tm62).antisense | CACTCATGGTTATGGCAGCACTGC |
| DGEK_Lxyl D133A fw | CATAATTCCGCCGGAGAAAAAGAAGCAGC |
| DGEK_Lxyl D133A rv | TTTTTCTCCGGCGGAATTATGAATATAAC |
| gfp-uDF-DGEKG-Lxyl-8953-fw | TTTGTATAGTTCATCCATGCCATG |
| GFP238 24rv | ATGGATGAACTATACAAAGCATGTTTTTTTTGTACGGCAGTA |
| Lxyl-DGEKE-F124-V125A-rv | TATATAAGCCGGATGGCCGTGATACGT |
| Lxyl-DGEKE-F124A-fw | CATCCGGCTTATATATCCCTCCCGC |
| Lxyl-DGEKE-H130A-fw | GTGATACGTTATATTGCCAATTCCGACGGA |
| Lxyl-DGEKE-H130A-rv | AATATAACGTATCACAAACATCCGGCTTATATATCCCT |
| Lxyl-DGEKE-I120A-S121A-rv | ATATCCCTCCCGCATAAGAACGGTG |
| Lxyl-DGEKE-I126A-fw | AGCCGGATGTTTGTGGCCCGTTATATT |
| Lxyl-DGEKE-I126A-R127A-rv | CACAAACATCCGGCTTATATATCCCTCCCG |
| Lxyl-DGEKE-I129A-fw | TTTGTGATACGTTATGCCCATAATTCCGACG |
| Lxyl-DGEKE-ktSD 31-fw | TAACTTTAAGAAGGAGGGAGATATACCAATGGAAACCAGGAGGAAAAAATCTGGG |
| Lxyl-DGEKE-M123A-fw | GATATATAAGCCGGGCCTTTGTGATACGTT |
| Lxyl-DGEKE-R122A-fw | GAGGGATATATAAGCGCCATGTTTGTGATACGT |
| Lxyl-DGEKE-R122A-M123A-rv | GCTTATATATCCCTCCCGCATAAGAACGG |
| Lxyl-DGEKE-R127A-fw | CGGATGTTTGTGATAGCCTATATTCATAATTCCGACG |
| Lxyl-DGEKE-S121A-fw | CGGGAGGGATATATAGCCCGGATGT |
| Lxyl-DGEKE-S132A-fw | CGTTATATTCATAATGCCGACGGAGAAAAAGAAG |
| Lxyl-DGEKE-S132A-rv | ATTATGAATATAACGTATCACAAACATCCGGCTT |
| Lxyl-DGEKE-ss6-25aa-fw2 | ACACAACTGAATACGGTTGATGTG |
| Lxyl-DGEKE-ss6-25aa-rv2 | CAGTTGTGTCCTCCTGGTTTCCATTGGTAT |
| Lxyl-DGEKE-V125A-fw | ATAAGCCGGATGTTTGCCATACGTTATATTCAT |
| Lxyl-DGEKE-Y128A-fw | ATGTTTGTGATACGTGCCATTCATAATTCCGA |
| Lxyl-DGEKE-Y128A-I129A-rv | ACGTATCACAAACATCCGGCTTATATATCCCT |
| myc24fw | GAACAAAAACTCATCTCAGAAGAG |
| PmiifM-ktSD 24-rv | CATTGGTATATCTCCCTCCTTCTTAAAGTTAATTTTACTATATGTACAAGCT |
| uDF-DGEKG-Lxyl-9302-myc-rv | TGAGATGAGTTTTTGTTCATTTTCAATCTGCCATAGCCTGCC |
| a63-Lxyl-98ins-fw | CTTAATTATATATGTTTAAAACACATTCTTCTTATTT |
| a64-Lxyl-K97-rv | TTTTAAACATATATAATTAAGGATTACGCTTTTTAG |
| a65-Lxyl-FS15-fw | ATTATGGTCACTTCGCACGTTCTTATGCGGGAGGGA |
| a66-Lxyl-FS15-rv | GTGCGAAGTGACCATAATGCAAAC |
| a67-Lxyl-FS20-fw | CACCGTTCTTATGCGGGAGGATATATAAGCCGG |
| a68-Lxyl-FS20-rv | TCCCGCATAAGAACGGTGCGAAGT |
| a69-Lxyl-FS25-fw | GGAGGGATATATAAGCCGATGTTTGTGATACGT |
| a70-Lxyl-FS25-rv | CGGCTTATATATCCCTCCCGCATA |
| a71-Lxyl-FS30-fw | CCGGATGTTTGTGATACGTATATTCATAATTCC |
| a72-Lxyl-FS30-rv | CGTATCACAAACATCCGGCTTATA |
| b27-mifM91-fw | GGTTCTCTTCTTTTATAAACCGCATTT |
| b36-DGMK-mifM91-rv | TTATAAAAGAAGAGAACCTCATTTCATCCCGTCTTTTTTATGAAG |
| b37-DGMK-D136A-mifM91-rv | TTATAAAAGAAGAGAACCTCATTTCATCCCGGCTTTTTTATG |
| b42-Lxyl_DGEKE-mifM91-rv | TTATAAAAGAAGAGAACCTTCTTTTTCTCCGTCGGAATTATG |
| b43-Lxyl_AGEKE-mifM91-rv | TTATAAAAGAAGAGAACCTTCTTTTTCTCCGGCGGAATTATG |
| b44-Lxyl_DGEK-ST-mifM91-rv | TTATAAAAGAAGAGAACCTTATTTTTCTCCGTCGGAATTATG |
| b45-Lxyl_AGEK-ST-mifM91-rv | TTATAAAAGAAGAGAACCTTATTTTTCTCCGGCGGAATTATG |
| b86-Lxyl_AGMK-ST-mifM91-rv | TTATAAAAGAAGAGAACCTTATTTCATTCCGGCGGAATTATG |
| b89-DGMK_DGEKE-mifM91-rv | TTATAAAAGAAGAGAACCTTCTTTTTCCCCGTCTTTTTTATGAAG |
| b90-DGMK_AGEKE-mifM91-rv | TTATAAAAGAAGAGAACCTTCTTTTTCCCCGGCTTTTTTATG |
| b93-DGEK-FY-fw | CATAATTCCGACGGAGAAAAATWCGCAGCAGGCAGGCTA |
| b94-DGEK-IR-fw | CATAATTCCGACGGAGAAAAACKAGCAGCAGGCAGGCTA |
| b95-DGEK-DG-fw | CATAATTCCGACGGAGAAAAAGMCGCAGCAGGCAGGCTA |
| b96-DGEKE-136-rv | TTTTTCTCCGTCGGAATTATGAATATAACG |
| gfp238-fw | GGCATGGATGAACTATACAAA |
| gfp238-rv | TTTGTATAGTTCATCCATGCC |
| 5_uD_Lind_DGEKG fw | CAGGTACATTCATGATTCAGACGGTGAAAAAGGTGTAAGGGCAGGAGTTTTAAAGGTAAGTGAACAAAAACTCATCTCAGAAGAG |
| 5_uD_Lind_DGEKG rv | TCTGAATCATGAATGTACCTGATCAGATATTCTCTGCCGGTATACCCTTCCTTCATCAGCACGGTCAGGGCAATGGCCTGGAAAG |
| DGEKG_Lind D133A fw | TATCTGATCAGGTACATTCATGATTCAGCCGGTGAAAAAGGTGTAAGGGC |
| DGEKG_Lind H130 rv | ATGAATGTACCTGATCAGATATTC |
| DGEK_Lxyl N131A fw | TATATTCATGCATCCGACGGAGAAAAAGA |
| DGEK_Lxyl N131A rv | TCCGTCGGATGCATGAATATAACGTATCA |
| DGEK_Lxyl D133A fw | CATAATTCCGCCGGAGAAAAAGAACGACG |
| DGEK_Lxyl D133A rv | TTTTTCTCCGGCGGAATTATGAATATAAC |
| DGEK_Lxyl G134A fw | AATTCCGACGCAGAAAAAGAAGCAGCAGG |
| DGEK_Lxyl G134A rv | TTCTTTTTCTGCGTCGGAATTATGAATAT |
| DGEK_Lxyl E135A fw | TCCGACGGAGCAAAAGAAGCAGCAGGCAG |
| DGEK_Lxyl E135A rv | TGCTTCTTTTGCTCCGTCGGAATTATGAA |
| DGEK_Lxyl K136A fw | GACGGAGAAGCAGAAGCAGCAGGCAGGCT |
| DGEK_Lxyl K136A rv | TGCTGCTTCTGCTTCTCCGTCGGAATTAT |
| DGEK_Lxyl E137A fw | GGAGAAAAAGCAGCAGCAGGCAGGCTATG |
| DGEK_Lxyl E137A rv | GCCTGCTGCTGCTTTTTCTCCGTCGGAAT |
| Lxyl-DGEKE-mifM SD-L1-M-fw | GAAGCTAGGAGGAGGATGTGATGGAAACCAGGAGGAAAAAATCTGGG |
| Lxyl-DGEKE-mifM SD-L1-M-rv | CATCCTCCTCCTAGCTTCATTTTACTATATGTACAAGCTGTTGTC |

**Supplementary Table 4: Degenerate primers for DMS**

| Primer name | DNA sequence (5' - 3') |
| --- | --- |
| Lxyl-DGEKE-G118-NNK-fw | ACCGTTCTTATGCGGGAGNNKTATATAAGCCGGATG |
| Lxyl-DGEKE-Y119-NNK-fw | GTTCTTATGCGGGAGGGANNKATAAGCCGGATGTTT |
| Lxyl-DGEKE-I120-NNK-fw | CTTATGCGGGAGGGATATNNKAGCCGGATGTTTGTG |
| Lxyl-DGEKE-S121-NNK-fw | ATGCGGGAGGGATATATANNKCGGATGTTTGTGATA |
| Lxyl-DGEKE-R122-NNK-fw | CGGGAGGGATATATAAGCNNKATGTTTGTGATACGT |
| Lxyl-DGEKE-M123-NNK-fw | GAGGGATATATAAGCCGGNNKTTTGTGATACGTTAT |
| Lxyl-DGEKE-F124-NNK-fw | GGATATATAAGCCGGATGNNKGTGATACGTTATATT |
| Lxyl-DGEKE-V125-NNK-fw | TATATAAGCCGGATGTTTNNKATACGTTATATTCAT |
| Lxyl-DGEKE-I126-NNK-fw | ATAAGCCGGATGTTTGTGNNKCGTTATATTCATAAT |
| Lxyl-DGEKE-R127-NNK-fw | AGCCGGATGTTTGTGATANNKTATATTCATAATTCC |
| Lxyl-DGEKE-Y128-NNK-fw | CGGATGTTTGTGATACGTNNKATTTCATAATTCCGAC |
| Lxyl-DGEKE-I129-NNK-fw | ATGTTTGTGATACGTTATNNKCATAATTCGACGGA |
| Lxyl-DGEKE-H130-NNK-fw | TTTGTGATACGTTATATTNNKAATTCGACGGAGAA |
| Lxyl-DGEKE-N131-NNK-fw | GTGATACGTTATATTCATNNKTCCGACGGAGAAAAA |
| Lxyl-DGEKE-S132-NNK-fw | ATACGTTATATTCATAATNNKGACGGAGAAAAAGAA |
| Lxyl-DGEKE-D133-NNK-fw | CGTTATATTCATAATTCNNKGGAGAAAAAGAAGCA |
| Lxyl-DGEKE-G134-NNK-fw | TATATTCATAATTCGACNNKGAAAAAGAAGCAGCA |
| Lxyl-DGEKE-E135-NNK-fw | ATTCATAATTCGACGGANNKAAAGAAGCAGCAGGC |
| Lxyl-DGEKE-K136-NNK-fw | CATAATTCGACGGAGAANNKGAAGCAGCAGGCAGG |
| Lxyl-DGEKE-E137-NNK-fw | AATTCGACGGAGAAAAANNKGCAGCAGGCAGGCTA |
| Lxyl-DGEKE-E117-rv | CTCCCGCATAAGAACGGTGCGAAGTGACCA |
| Lxyl-DGEKE-Y119-rv | ATATCCCTCCCGCATAAGAACGGTGCGAAG |
| Lxyl-DGEKE-S121-rv | GCTTATATATCCCTCCCGCATAAGAACGGT |
| Lxyl-DGEKE-M123-rv | CATCCGGCTTATATATCCCTCCCGCATAAG |
| Lxyl-DGEKE-V125-rv | CACAAACATCCGGCTTATATATCCCTCCCG |
| Lxyl-DGEKE-R127-rv | ACGTATCACAAACATCCGGCTTATATATCC |
| Lxyl-DGEKE-I129-rv | AATATAACGTATCACAAACATCCGGCTTAT |
| Lxyl-DGEKE-N131-rv | ATTATGAATATAACGTATCACAAACATCCG |
| Lxyl-DGEKE-D133-rv | GTCGGAATTATGAATATAACGTATCACAAA |
| Lxyl-DGEKE-E135-rv | TTCTCCGTCCGGAATTATGAATATAACGTAT |

**Supplementary Table 5: Primers used for preparation of NGS library**

| Primer name | Sequence (5'-3') |
| --- | --- |
| P5-UDI0001-Rd1-GGT-Lxyl_AP-fw | AATGATACGGCGACCACCGAGATCTACACAGCGCTAGACACTCTTTCCCTACACGACGCTCT<br>TCCGATCTNNNNNNGGTTTTCTTAATATTGTAAGCCGC |
| P7-UDI0001-Rd2-ACC-FLAG2-8-rv | CAAGCAGAAGACGGCATACGAGATAACCGCGGGTGACTGGAGTTCAGACGTGTGCTCTTCC<br>GATCTNNNNNNNACCTTTGTCGTCGTCGTCCTTTATA |
| P7-UDI0001-Rd2-AGC-FLAG2-8-rv | CAAGCAGAAGACGGCATACGAGATAACCGCGGGTGACTGGAGTTCAGACGTGTGCTCTTCC<br>GATCTNNNNNNNAGCTTTGTCGTCGTCGTCCTTTATA |
| P7-UDI0001-Rd2-AGG-FLAG2-8-rv | CAAGCAGAAGACGGCATACGAGATAACCGCGGGTGACTGGAGTTCAGACGTGTGCTCTTCC<br>GATCTNNNNNNNAGGTTTGTCTCGTCGTCGTCCTTTATA |
| P7-UDI0001-Rd2-CTC-FLAG2-8-rv | CAAGCAGAAGACGGCATACGAGATAACCGCGGGTGACTGGAGTTCAGACGTGTGCTCTTCC<br>GATCTNNNNNNNCTCTTTGTCGTCGTCGTCCTTTATA |
| P7-UDI0001-Rd2-CTG-FLAG2-8-rv | CAAGCAGAAGACGGCATACGAGATAACCGCGGGTGACTGGAGTTCAGACGTGTGCTCTTCC<br>GATCTNNNNNNNCTGTTTGTCTCGTCGTCGTCCTTTATA |
| P7-UDI0001-Rd2-TGC-FLAG2-8-rv | CAAGCAGAAGACGGCATACGAGATAACCGCGGGTGACTGGAGTTCAGACGTGTGCTCTTCC<br>GATCTNNNNNNNTGCTTTGTCTCGTCGTCGTCCTTTATA |
| P7-UDI0001-Rd2-TTA-FLAG2-8-rv | CAAGCAGAAGACGGCATACGAGATAACCGCGGGTGACTGGAGTTCAGACGTGTGCTCTTCC<br>GATCTNNNNNNNTTATTTGTCTCGTCGTCGTCCTTTATA |
| P7-UDI0001-Rd2-TTG-FLAG2-8-rv | CAAGCAGAAGACGGCATACGAGATAACCGCGGGTGACTGGAGTTCAGACGTGTGCTCTTCC<br>GATCTNNNNNNNTTGTGTCGTCGTCGTCCTTTATA |

##### **Supplementary Movie Legend**

Cryo-EM map of LudP-SRC stalling during elongation, including a section that reveals the quality of the density (transparent surfaces) for the nascent chain (mint), P-tRNA (purple) and the A-tRNA (salmon) together with the molecular model. The final part of the movie shows a clash of D133 of LudP (mint) with U2506 (dark grey)(PDB ID 8CVK)<sup>6</sup> from the 23SrRNA in the pre-attack state. Therefore, U2506 (grey) is displaced into the A-site of the PTC which causes elongation and termination arrest through steric hinderance of A-tRNA (brown)(PDB ID 8CVK)<sup>6</sup> and RF (green)(PDB ID 9MTP)<sup>7</sup> accommodation in the A-site of the PTC.
